# The dominant role of post-transcriptional regulation in *Yarrowia lipolytica* to repeated O_2_ limitations

**DOI:** 10.1101/2023.07.13.548809

**Authors:** Abraham Antonius Johannes Kerssemakers, Mariam Nickseresht Funder, Süleyman Øzmerih, Suresh Sudarsan

## Abstract

Rational scale-up strategies to accelerate bioprocess development, require sound knowledge of cellular behaviour under industrial conditions. In this study, the strictly aerobic yeast *Yarrowia lipolytica* is exposed to repeated oxygen limitations, approximated from a large-scale cultivation. A data-driven multi-omics strategy is deployed to elucidate its transcriptomic, proteomic, and metabolic response. Throughout a single perturbation, metabolite and protein levels showed dynamic profiles while they returned to steady state values when aerobic conditions were restored. After repeated oscillations, significant cellular rearrangements were found, with a special focus on central carbon metabolism, oxidative phosphorylation, lipid, and amino acid biosynthesis. Most notably, metabolite levels as well as the catabolic reduction charge are maintained at higher concentrations. Moreover, proteins involved in NADPH-consuming anabolic pathways showed an increased abundance, which is suggested to be compensated for through an increased pentose-phosphate pathway activity. Although dynamics were found on all three omics levels, the proteomic and metabolic changes were in most instances not supported by strong transcriptional changes. Thus, this work suggests that the response of *Y. lipolytica* to (repeated) oxygen oscillations is strongly regulated by post-transcriptional mechanisms. These findings provide novel insights into potential cellular regulation on an industrial scale, thereby facilitating a more efficient bioprocess development through mitigating any undesired behaviour.

**Key points:**

- Dynamic response of *Yarrowia lipolytica* to industrial oxygen profiles.
- New metabolic steady states are found after exposure to repeated oxygen oscillations.
- A multi-omics strategy elucidates the importance of post-transcriptional mechanisms.

## Introduction

Microbial catalysis has the potential to play a crucial role in the transition to a cleaner and more renewable way of meeting our needs (UN 2015; EU 2019); through the production of a wide selection of compounds, ranging from food products to energy and chemicals (Mengal et al. 2018). For several decades already, academia and industry have developed a wide variety of bioproduction pathways, making use of different feedstocks, fermentation setups, and microbial hosts. Still, one of the main challenges in industrial bioprocess development is the successful transfer of processes from the laboratory to industrial-sized vessels while maintaining desired microbial performance (Straathof et al. 2019). Although microbial behavior in laboratory setups is relatively well understood, this so-called scale-up is often complicated through undesired industrial characteristics (Takors 2012). One of the main contributors to this challenge is the difference in bioreactor size, which leads to altered physical/chemical characteristics on a larger scale. Whereas laboratory setups are relatively homogenous and can be tightly controlled, larger volumes are accompanied by increased heterogeneities (Wehrs et al. 2019). As a result, production strains might be exposed to suboptimal conditions, thereby not fulfilling their maximum potential regarding titers, rates, and yields.

Thus, to have a more efficient scale-up, it is of great importance to have an understanding of the heterogeneities that industrial vessels bring (Nadal-Rey et al. 2021b). Great efforts are made in this field through various efforts such as computational fluid dynamics and the related compartmental modelling (Tajsoleiman et al. 2019; Wang et al. 2020). These fields output a description of crucial concepts such as liquid and gas mixing. When combining these results with microbial kinetics, an accurate approximation can be made of the conditions encountered in an industrial vessel (Haringa et al. 2018; Nadal-Rey et al. 2021a). By doing so, a reference framework is designed in which microbes can be tested in the laboratory. This approach is the so-called scale-down, which allows researchers to bring industrial heterogeneities into the laboratory and thus have a more predictive setup for bioprocess development (Noorman 2011).

In most cases, the setup is one in which environmental conditions continuously change over time. This raises the question as to how microbes respond to these continuously oscillating conditions. Microbial systems will adjust their machinery based on the environment to which they are exposed (Lelli et al. 2012). Consequently, oscillating reactor conditions will result in dynamic microbial behaviour (Lee et al. 2002). It’s crucial to achieve a better understanding of these dynamics as that is often the cause for the changes in productivity that make scale-up so challenging. A constant rearrangement of cells’ transcriptomic, proteomic, and/or metabolic landscape could result in a decreased efficiency (Eigenstetter and Takors 2017).

Often, it can be challenging to change the physical properties of an industrial vessel, as they are intrinsic challenges that come with the increase in size. As a result, solutions will also have to be found in the microbial domain. Thus, researchers aim to identify and create the so-called *fermenterphile* geno- and phenotype (Straathof et al. 2019). These strains show robustness to industrial conditions and thus have an unaltered performance when brought to a commercial scale.

In this study, there is a special focus on *Yarrowia lipolytica*. This strictly aerobic, oleaginous yeast is gaining in popularity due to its potential to grow on a wide range of substrates, relevant products, general robustness and increasingly well understood genomic makeup (Coelho et al. 2010). In relation to scale-up efforts, robustness of a strain is a great asset. Being a strict aerobe however, oxygen availability is of main concern when growing in larger volumes and higher cell densities. To aid in the development of industrial *Y. lipolytica* based bioprocesses, this study aims to deepen the understanding of its cellular makeup and control. A relevant scale-down study is designed, specifically targeting continuous oxygen oscillations. The transcriptome, proteome and metabolome are studied and integrated in a multi-omics approach to better elucidate the changes in *Y. lipolytica* in response to these industrial stresses.

## Material and methods

### Cultivation conditions

The wild-type *Yarrowia lipolytica* W29 (ATCC 20460) was used in this research (Nicaud 2012).

Pre-cultures were prepared in 250 mL shake flasks by using 250 µL stock culture (25% glycerol, -80 °C) to inoculate 25 mL Delft media (20 g/L glucose, 14.4 g/L KH_2_PO_4_, 0.5 g/L MgSO_4_, 7.5 g/L (NH_4_)2SO_4_, 0.1mg/L biotin, 0.4 mg/L 4-aminobenzoic acid, 2 mg/L nicotinic acid, 2 mg/L calcium pantothenate, 2 mg/L pyri-doxine, 2 mg/L thiamine HCl, 50 mg/L myo-inositol, 4.5 mg/L CaCl_2_·2H_2_O, 4.5 mg/L ZnSO_4_·7H_2_O, 3 mg/L FeSO_4_·7H_2_O, 1mg/L boric acid, 1 mg/L MnCl_2_·4H_2_O, 0.4 mg/L Na_2_MoO_4_·2H_2_O, 0.3 mg/L CoCl_2_·6H_2_O, 0.1 mg/L CuSO_4_·5H_2_O, 0.1 mg/L KI and 15 mg/L EDTA) and left to grow overnight at 30 °C and 225 RPM.

Continuous cultivations were performed in a 1L Biostat Q Plus vessel (Sartorious, France) with a working volume of 0.5 L. Dissolved oxygen (DO) was measured with the VisiFerm DO 160 probe (Hamilton Company, US) and controlled by adjusting the flowrate of air, and by mixing with pure O_2_ or N_2_. A minimum flow rate of 0.6 vvm was used to ensure that sufficient airflow was supplied to the PrimaBT Mass Spectrometer (Thermo Scientific, USA). Stirring was done with a single Rushton impeller at a constant speed of 500 RPM to avoid increased liquid levels due to increased agitation. pH was monitored with the EasyFerm Plus PHI k8 160 probe (Hamilton Company, US) and maintained at pH 6 by automatic addition of 5M NaOH using the built-in PID control loop. The temperature was set at 30 °C and was controlled automatically. Feed rates were controlled by external pumps that were calibrated prior to use. Media was designed based on the Delft composition with a glucose concentration of 10 g/L and the addition of 1 mL/L of Antifoam 204 (Merck, Germany).

The bioreactors were inoculated at an OD_600_ of 0.1. After completion of the batch-phase, feeding and harvest pumps were activated to maintain a constant dilution rate. Eight residence times passed before reaching the steady state. Then, for the first stimulus-response experiment, aeration was fully interrupted until cellular activity brought the DO to 0%. After 2 minutes of anaerobic conditions, aeration was turned on again (0.6 vvm air) to restore the original DO value of 40%. This first oscillation was followed by a 10 hour profile in which the bioreactor was alternatingly sparged with 0.2 vvm N_2_ for 2.5 minutes followed by 0.2 vvm O_2_ for 2.5 minutes.

### Sampling for biomass determination, transcriptome, proteome, and metabolome

The cell dry weight (CDW) was determined by drying known sample volumes after washing the cell pellet twice with a 0.9% NaCl solution. The samples were placed in pre-dried and weighed Eppendorf tubes and dried in an oven at 80 °C until a constant weight was reached. For transcriptome and proteome samples, 1 mL of broth was sampled in pre-cooled Eppendorf tubes and centrifuged directly for one minute at 4000 g and 2 °C. The supernatant was discarded and the remaining cell pellet was snap-frozen in liquid nitrogen. Metabolomic samples were taken through rapidly vacuum extracting (< 1s) 1 mL of broth into 9 mL of 80% (v/v) MeOH precooled at -40 °C (Canelas et al. 2008). After sampling, the tubes were directly vortexed and weights were measured before and after sample extraction to determine the exact amount of sample taken. Then, sample tubes were spun down at 4000 g for 5 minutes at -20 °C. The supernatant was discarded and the remaining cell pellet was washed in 5 mL of precooled 80% (v/v) MeOH and centrifuged and decanted as before. The pellets were then extracted through the boiling ethanol method (Canelas et al. 2009). 5 mL boiling 75% (v/v) ethanol was added to the tubes which were then vortexed and placed in a water bath at 95 °C for three minutes. Sample tubes were then rapidly cooled down and stored at -80 °C until further analysis.

### Analysis of intracellular metabolome

The ethanol fraction was evaporated off the cell pellet. Samples were reconstituted in water (HPLC grade H2O) and analyzed on an UHPLC-ESI-QTrap-MS (UHPLC LC20/30 series, Schimadzu and QTrap, Sciex) using a XSelect HSS T3 XP column (2.1x150mm, particle size 2.5 µm, Waters) as previously described (McCloskey et al. 2015). Data was acquired using the scheduled MRM algorithm in Analyst Software (Sciex). Post-run data analysis was performed in SciexOS Software (Sciex). Compounds were quantified using ^13^C metabolically labeled internal standards from *E. coli*. Linear regressions for quantification were based on peak height ratios. The lower limit of quantification was defined at a peak height greater than 1e^3^ ions counts and signal to noise ratio greater than 15 (McCloskey et al. 2015).

The following binary solvent gradient (A: 10mM Tributylamine, 10mM Acetic Acid, 5%[v/v] MeOH, 2%[v/v] Isopropanol in water; and B: Isopropanol) was applied: 0 to 5 min, isocratic 100% A; 5 to 9 min, linear ramp to 98% A, 2% B; 9 to 9.5 min, linear ramp to 94% A, 6% B; 9.5 to 11.5 min, isocratic 94% A, 6% B; 11.5 to 12 min, linear ramp to 89% A, 11% B; 12 to 13.5 min, isocratic 89% A, 11% B; 13.5 to 15.5 min, linear ramp to 72% A, 28% B; 15.5 to 16.5 min, linear ramp to 47% A, 53% B; 16.5 to 22.5 min, isocratic 47% A, 53% B; 22.5 to 23 min, linear ramp to 100% A; 23 to 27 min, isocratic 100% A. The flow rate was 0.4 ml min^−1^ from 0 to 15.5 min, 0.15 ml min^−1^ from 15.5 to 23 min and 0.4 ml min^−1^ from 23 to 27 min. The injection volume was 10 μl. MS data was acquired from 0.4 to 23 min (McCloskey et al. 2015).

To calculate the energy, catabolic and anabolic reduction charges, the following equations were used (Ball and Atkinson 1975; Nielsen et al. 2003):

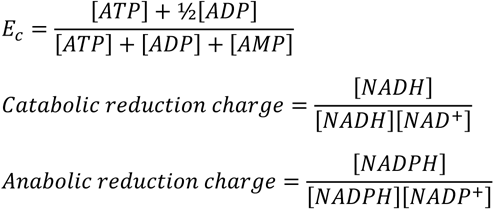

### Analysis of cellular proteome

Peptides were loaded onto a 2 cm C18 trap column (ThermoFisher 164946), connected in-line to a 15 cm C18 reverse-phase analytical column (Thermo EasySpray ES904) using 100% Buffer A (0.1% Formic acid in water) at 750 bar, using the Thermo EasyLC 1200 HPLC system, and the column oven operating at 30 °C. Peptides were eluted over a 70 min. gradient ranging from 10% to 60% of Buffer B (80% acetonitrile, 0.1% formic acid) at 250 nl/min, and the Orbitrap Exploris instrument (Thermo Fisher Scientific) was run in DIA mode with FAIMS ProTM Interface (ThermoFisher Scientific) with CV of -45 V. Full MS spectra were collected at a resolution of 120,000, with an AGC target of 300% or maximum injection time set to ‘auto’ and a scan range of 400–1000 m/z. The MS2 spectra were obtained in DIA mode in the orbitrap operating at a resolution of 60,000, with an AGC target 1000% or maximum injection time set to ‘auto’, a normalised HCD collision energy of 32. The isolation window was set to 6 m/z with a 1 m/z overlap and window placement on. Each DIA experiment covered a range of 200 m/z resulting in three DIA experiments (400-600 m/z, 600-800 m/z and 800-1000 m/z). Between the DIA experiments a full MS scan is performed. MS performance was verified for consistency by running complex cell lysate quality control standards, and chromatography was monitored to check for reproducibility.

The raw files were analyzed using SpectronautTM (version 16.2), spectra were matched against the Uniprot *Yarrowia lipolytica* database. Dynamic modifications were set as Oxidation (M) and Acetyl on protein N-termini. Cysteine carbamidomethyl was set as a static modification. All results were filtered to a 1% FDR, and protein quantitation done on the MS1 level. The data was normalized by RT dependent local regression model and protein groups were inferred by IDPicker (Callister et al. 2006).

### Analysis of transcriptome

RNA extraction, quality control, library preparation, and paired-end sequencing were performed by an external service partner (BGI Europe, Copenhagen, Denmark). The alignment was done using nf-core rnaseq, combining Star and Salmon to align and quantify the reads (Ewels et al. 2020). The QC method is a 2-step process. First, samples were removed based on the nf-core rnaseq multiqc report. Then gene expression profiles of the samples were clustered and analysed to remove outlier clusters within which the samples were not biologically related. Here, the correlation between all the sample replicates and samples was computed at random. Any sample with a replicate correlation < 0.9 was rejected.

The resulting transcript per million (TPM) values allowed for an expression analysis through DESeq2 and the results were used for a principal component analysis (Love et al. 2014). A differential gene expression analysis was performed based on the TPM log2 difference against the steady state condition. Log2 differences lower than −1 indicated a downregulation, whereas values higher than 1 were considered upregulated.

## Results

### Metabolic response of *Y. lipolytica* to oscillations in dissolved O2

As *Y. lipolytica* is a strict aerobe, fluctuating oxygen availabilities are suspected to have an effect on its performance. Thus, from the perspective of developing a *fermenterphile* strain, it is crucial to gain a better understanding of these dynamics. In this study, this was achieved through a continuous culture with oxygen oscillations. The continuous culture was achieved at a growth rate of 0.1 h^-1^, biomass concentration of 6.5 g_CDW_ L^-1,^ and a DO concentration of 40%. This steady state is considered the reference state after which an oxygen oscillation profile was introduced in the bioreactor. Dissolved oxygen values varied between 0 and 40%, mimicking an industrially relevant heterogeneity profile (Kerssemakers et al. 2023). The first oscillation acts as a stimulus-response setup and has a duration of 10 minutes before restoring the original DO value. Consecutive oscillations occur over a window of 5 minutes, which more closely resembles but is slightly higher than the calculated liquid mixing time of industrially sized fermenters (Gaugler et al. 2022; Nadal-Rey et al. 2022). The culture was sampled over the course of the first oscillation and during an aerobic and anaerobic phase at 10 hours into the oscillating profile. The resulting profile (Figure 1A) allows for the analysis of both short and long-term effects of (repeated) oxygen limitation.

**Figure 1.**
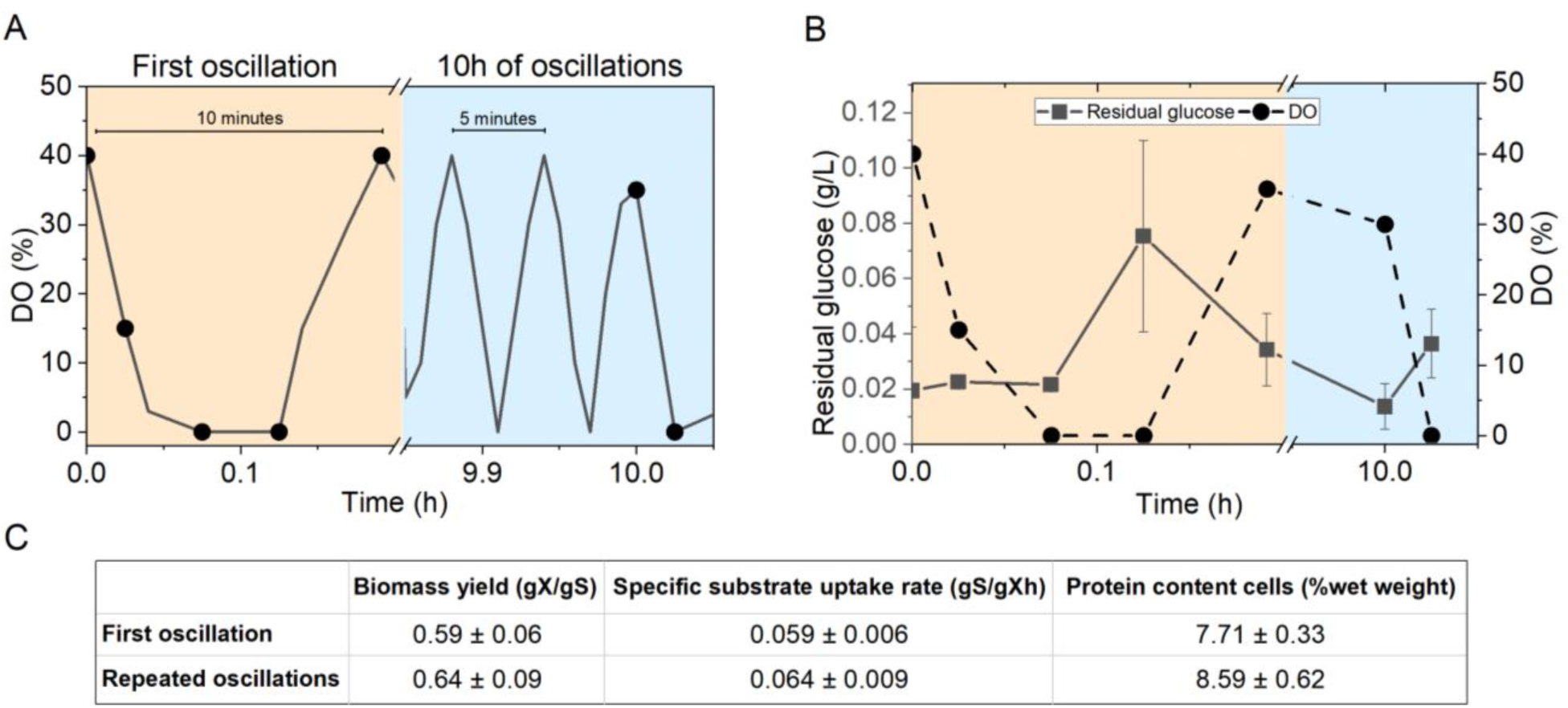
Overview of experimental setup, sampling times for metabolomics and kinetic parameters. A. Schematic overview of the experimental setup and metabolomics sampling time points. The continuous culture is brought to a steady state at a dissolved oxygen concentration of 40% and biomass concentration of 6.5 g_CDW_/L. After 80 hours (8 residence times) the oxygen oscillation profile is started (time = 0). After this initial oscillation, the profile continues for 10 hours with oxygen concentrations oscillating between 0 and 40% over a period of 5 minutes. Black dots indicate the sampling time points for metabolomics. B. Residual glucose concentration as a result of changing dissolved oxygen concentrations. C. Biomass yields, specific substrate uptake rates, and extracted protein concentrations at the start of the first oscillation and during the aerobic phase 10 hours into the repeated oscillations.

As *Y. lipolytica* is a strictly aerobic organism, oxygen is required for basic metabolic functioning and substrate utilization. Thus, under anaerobic conditions, it is expected that substrate will accumulate, which was confirmed through analysis of residual glucose concentrations (**Figure *1***B). While reaching anaerobic conditions, a spike in glucose concentrations is observed followed by complete consumption once aerobic conditions are restored. After repeated oscillations, there appears to be no substrate accumulation in the aerobic phase, while the anaerobic phase does again result in a small increase in residual glucose concentrations. This indicates that *Y. lipolytica* can metabolize all substrate supplied during the aerobic phase of the cycle. The biomass yields at the start of the first oscillation are slightly higher than during the aerobic phase after 10 hours of oscillations (**Figure *1***C). As all glucose is consumed, this results in higher specific substrate uptake rates. Moreover, the extracted protein concentrations are higher after repeated oscillations. Standard deviations, however, make it difficult to conclude on the statistical significance of the changes in these parameter values.

Key central metabolite concentrations were quantitatively determined for each sampling time point (Figure 2). The first measurement is taken during steady state and acts as the reference value, which allows for the study of dynamic behaviour of the metabolite pools. A comparison of central carbon metabolite steady state values to those reported in literature on *Y. lipolytica, Pichia pastoris* and *Saccharomyces cerevisiae* gives varying results (Table S1).

**Figure 2.**
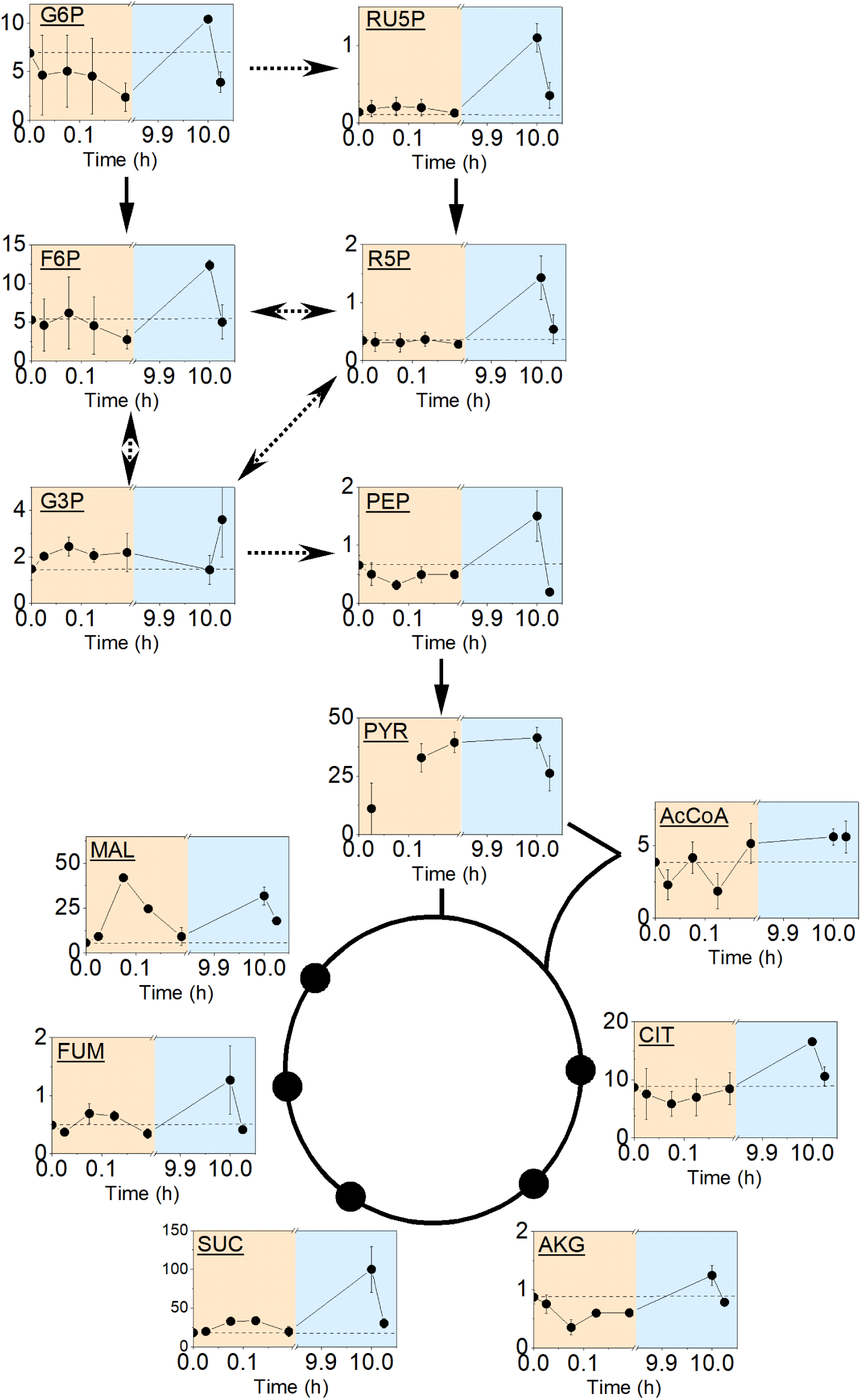
Dynamics of metabolites in central carbon metabolism to oscillating dissolved O_2_. In total, 13 central carbon metabolites were quantitatively measured over the course of an oxygen oscillation experiment. Orange boxes indicate the samples taken during the first oscillation and blue boxes represent the samples taken 10 hours into the oscillating profile. The first sample was taken during steady state conditions and therefore acts as the reference value to all other measurements (dotted line). All concentrations in µmol/g CDW asides from citrate which is shown in mmol/g CDW.

For the anaerobic phase in the first oscillation, a depletion of glucose 6-phosphate (G6P) and phosphoenolpyruvate (PEP) was observed, while glyceraldehyde 3-phosphate (G3P) and Pentose Phosphate Pathway (PPP) metabolite ribulose 5-phosphate (RU5P) show an accumulation. In the TCA cycle, a short-term depletion of citrate (CIT) and α-ketoglutarate (AKG) is seen, whereas succinate (SUC), fumarate (FUM) and malate (MAL) accumulate. Interestingly, this changing trend in the TCA cycle coincides with succinate dehydrogenase, an enzyme whose activity is closely linked to oxygen availability through the electron transport chain. After 2 minutes of anaerobic conditions, aeration is turned on again and aerobic conditions are restored. As oxygen becomes available, most metabolite concentrations again approximate their steady state values.

After repeated oscillations, the trends that were observed on the short term have altered. Now all metabolites, apart from G3P, show a depletion when transitioning into the anaerobic phase of the oscillation. Interestingly, many metabolites show a deviation from their steady state concentrations in the aerobic phase after being exposed to repeated oscillations. The difference is most noteworthy for the PPP metabolites, which show concentrations up to 8-fold higher than the steady state. Similarly, all TCA cycle intermediates show an increased concentration as compared to their steady state values.

Further, to better elucidate the effect of (repeated) oxygen limitation on *Y. lipolytica*, the energy and redox states of the cells have been assessed. Adenylate profiles were monitored and show an accumulation of AMP and ADP but depletion of ATP in the anaerobic phase of both the initial and repeated oscillating profile (Figure 3). Over the first oxygen cycle, AMP and ADP concentrations appear to return to their original steady state values. ATP on the contrary has a slight overshoot as the bioreactor returns to aerobic conditions. After repeated oscillations, all adenylates steady state values appear to be more strictly maintained as compared to the central carbon metabolites.

**Figure 3.**
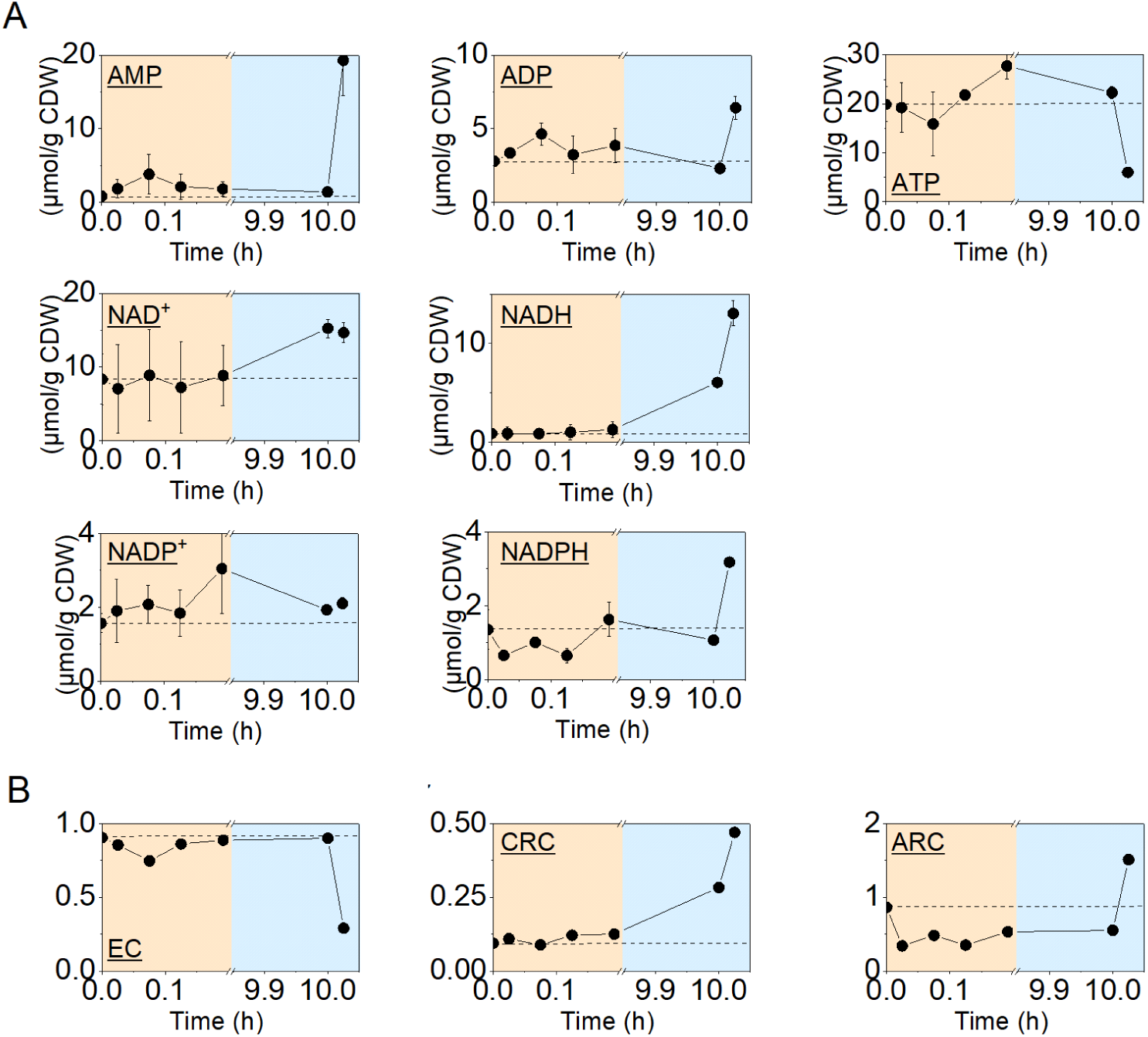
Energy and cofactor profiles. A. Dynamics of adenylates (AMP, ADP and ATP) and metabolite cofactors (NAD^+^, NADH, NADP^+^, NADPH in response to oscillations in dissolved O_2_. B. The energy charge (EC), catabolic reduction charge (CRC) and anabolic reduction charge (ARC) can be calculated based on these values. Dotted lines indicate the steady state values of the cells.

Based on these measured concentrations, the energy charge of the cells can be determined, which is an indicator of the amount of metabolically available energy. The calculated energy charge has a steady state value of 0.9 (**Figure *3***). The value decreases over the first oxygen oscillation to a minimum of 0.75, indicating a decreased energy availability. As oxygen is re-introduced into the system, the energy balance recovers to its original steady state value. After repeated oscillations, cells are able to maintain the energy charge of 0.9 in the aerobic part while values drop to 0.29 in the anaerobic phase. The rapid changes in the energy charge highlight the fast turnover rate of these compounds. When depleted of oxygen, ATP levels drop significantly in a matter of seconds to minutes. As the Gibbs free energy released by hydrolysis of ATP is used to drive biosynthetic pathways, this ultimately leaves the cells without energy for anabolic and catabolic functioning.

Analysis of redox couples NADPH/NADP^+^ and NADH/NAD^+^ provides additional fundamental insights into co-factor homeostasis in cells. NADP(H) is oxidized to NAD(P)^+^ with the release of two electrons, these can then be used to reduce other molecules in the cells. Over the initial oscillation, co-factor concentrations are relatively noisy and thus difficult to interpret. After repeated oscillations, however, an increase in NADH and NADPH was observed when transitioning into the anaerobic phase whereas NAD^+^ and NADP^+^ remained constant.

A set of parameters has been identified to adequately assess these redox states, namely the catabolic reduction charge and anabolic reduction charge. The steady state catabolic reduction charge is maintained at 0.1 and does not show significant changes over the course of the initial oscillation. NAD^+^ is a substrate in many catabolic reactions and its regeneration is largely dependent on electron transport complex I (NADH dehydrogenase). Thus, under anaerobic conditions, depletion of NAD^+^ and accumulation of NADH would be expected resulting in an increased reduction charge. Indeed, after repeated oscillations, the value rises from 0.28 under aerobic to 0.47 under anaerobic conditions. This also indicates a significant shift from the steady state value of 0.1 meaning that shifts in redox homeostasis have occurred.

The catabolic reduction charge is generally kept low, as NAD^+^ is a substrate and NADH a product in catabolic reactions. This contrasts with the anabolic reduction charge, which is generally kept at higher levels. Here NADPH is a substrate and NADP^+^ a product. Indeed, a steady state anabolic reduction charge of 0.8 was found in this study. After repeated oscillations, a slightly lower value is found under aerobic conditions. This indicates that the anabolic reduction homeostasis has been altered which supports the small changes observed in biomass yields. Once the system becomes anaerobic again, the anabolic reduction charge increases, which is the opposite of the behaviour observed under the short-term oxygen limitation. The main reason is a spike in NADPH levels, which could be related to the higher concentrations of PPP metabolites.

### Proteome and transcriptome response of *Y. lipolytica* to repeated O2 oscillations

Analysis of the proteome and transcriptome under oxygen-oscillating conditions provides an additional layer of depth to the analysis performed in this work (Figure 4A). Metabolic changes can be triggered by a variety of mechanisms and therefore it is important to differentiate between passive and active forms of metabolic regulation. In literature, this difference has also been described as hierarchical vs. metabolic regulation (Kuile and Westerhoff 2001). To address this question, a better understanding of the relation between proteome and transcriptome is required. As such, a Pearson correlation coefficient was calculated comparing the levels of transcription versus those of the corresponding proteins. Here a coefficient of 0.607 was found for the steady state conditions (Figure 4B). Over the course of the first oscillation, this coefficient gradually drops to 0.570 and returns to approximately 0.630 during the aerobic and anaerobic phase after repeated oscillations (Figure S1).

**Figure 4.**
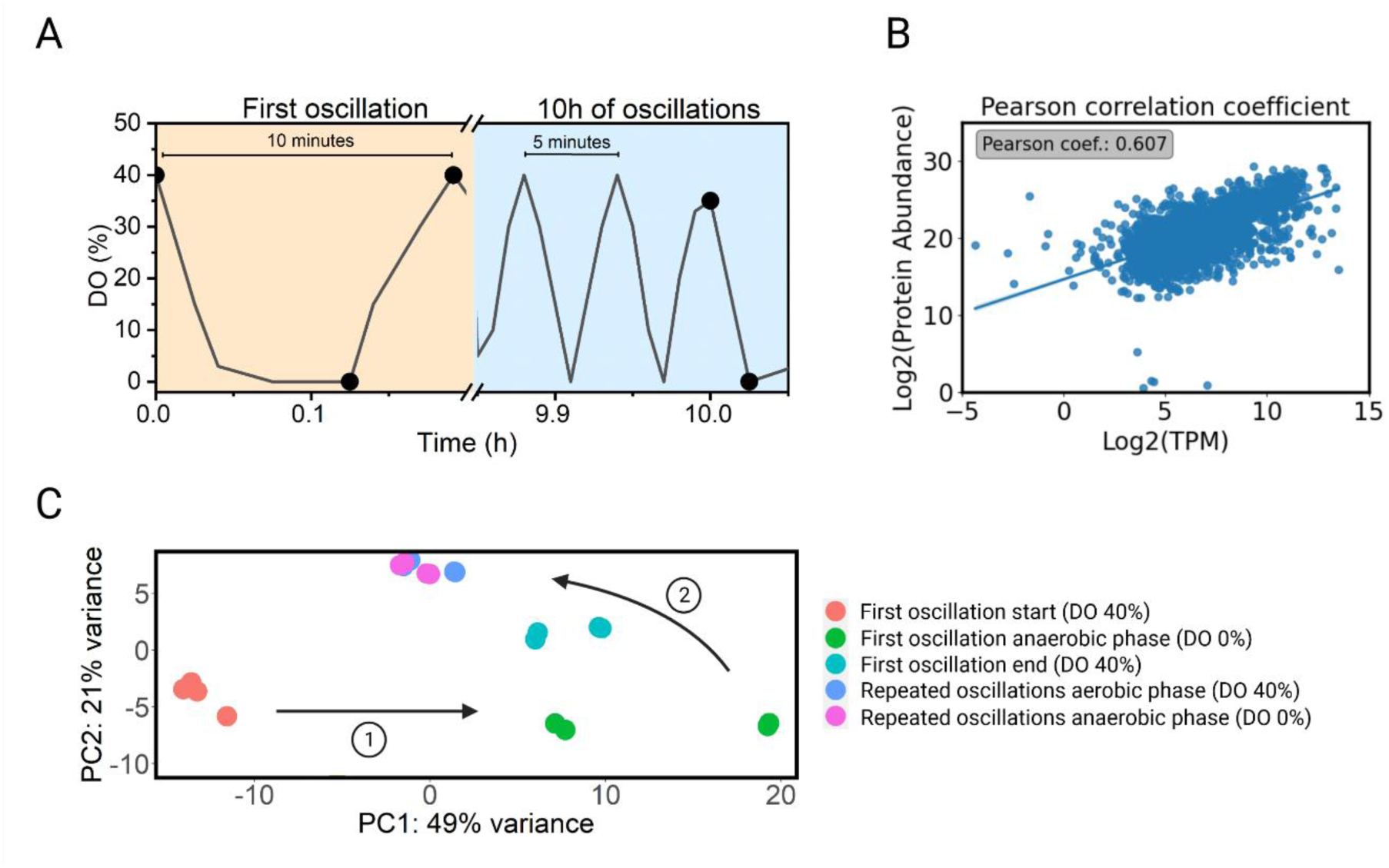
Global analysis of the proteome and transcriptome in *Y. lipolytica* exposed to O_2_ oscillations. A. Oxygen profile and sampling time points for proteomics and transcriptomics. B. Pearson correlation coefficient comparing protein abundance and the mRNA levels in TPM of the related genes. C. Principal Component Analysis of the transcriptomic data obtained from the oscillation experiment.

Then, to obtain a global overview of the effects of (repeated) oxygen limitation on *Y. lipolytica*, a principal component analysis was performed (Figure 4C). Principal component (PC) 1 and 2 explain a majority of the observed variance. As the initial oscillation progresses and the bioreactor becomes anaerobic, data points move across PC1. When aerobic conditions are reintroduced, datapoints move back in the direction of steady state conditions but do not yet fully reach the same state. After repeated oscillations the datapoints are clustered together, irrespective of whether they were taken during the aerobic or anaerobic phase. Interestingly, their location on the PC1-PC2 grid is different than that of the steady state condition, indicating that the transcriptomic landscape has changed.

### Catabolic shifts

Here, the central carbon metabolism has been split into five different pathways. For each pathway all proteins known to be present in *Y. lipolytica* were analysed (Figure 5A). The abundance of the protein during steady state conditions was taken as reference point and for all other conditions, a log2 fold change was calculated. As a result, proteins could either be more abundant (log2 fold change >1), less abundant (log2 fold change <-1) or unaltered (log 2fold change between -1 and 1).

**Figure 5.**
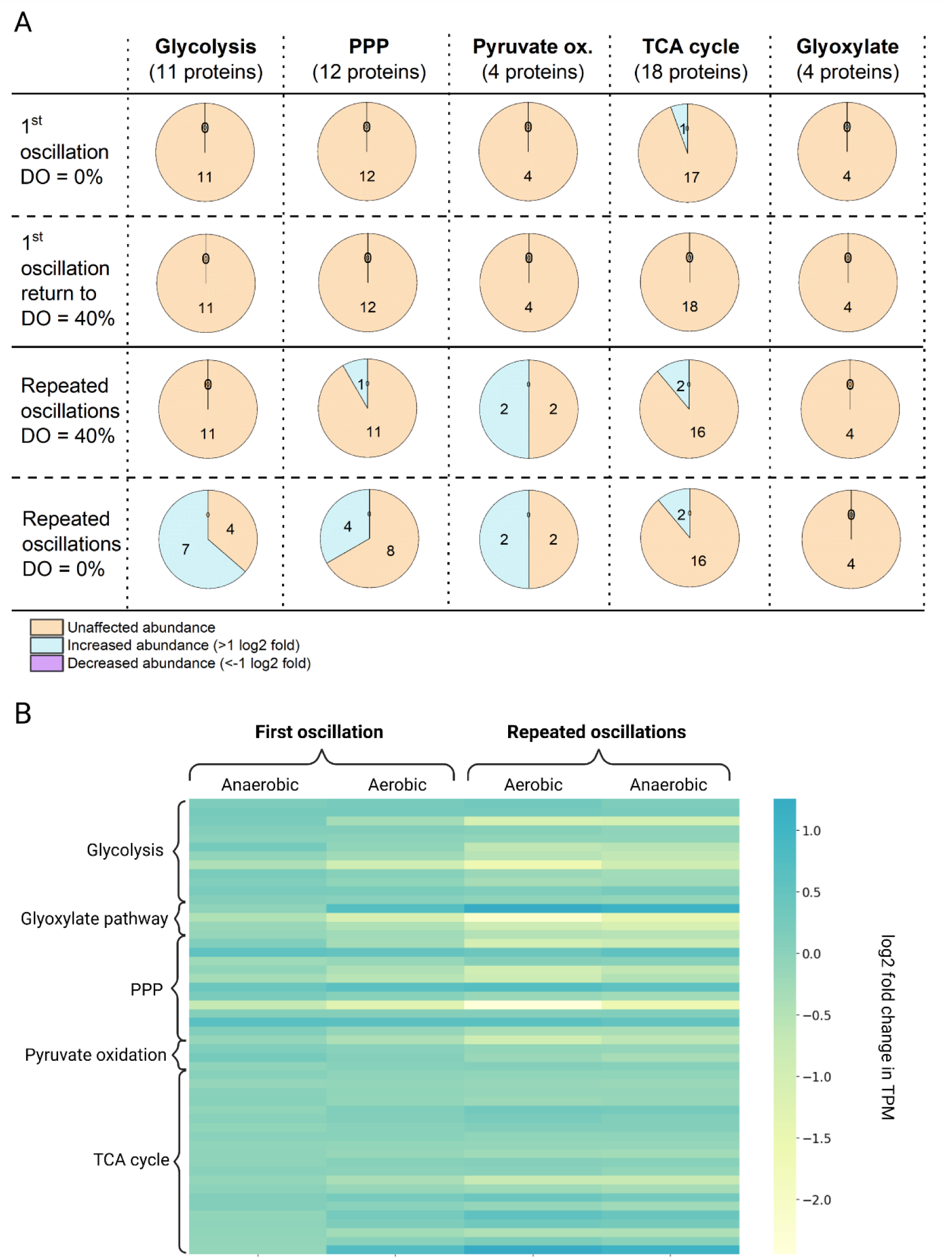
Proteomic and transcriptomic analysis of central carbon metabolism in response to oxygen oscillations. A. Protein abundance in central carbon metabolism pathways. Depending on the pathway that they’re involved in, proteins are divided into five different categories (Glycolysis, the Pentose Phosphate Pathway (PPP), Pyruvate oxidation, TCA cycle, and Glyoxylate pathway). Then over the course of the oscillation study, the log2 differences in protein abundance are calculated as compared to the reference steady state. Here the first row represents the protein abundance during the anaerobic phase of the first oscillation. The second row represents the end of the first perturbation when aerobic conditions are restored. The third row is the aerobic phase after repeated oscillations and the fourth row represents the anaerobic phase after repeated oscillations. A log2 fold change <-1 indicates are decreased abundance, >1 an increased abundance, and all values in between are considered not significant. B. log2 fold changes in TPM values of genes involved in central carbon metabolism.

Over the course of the first oscillation only succinate dehydrogenase, changes in abundance. This protein is involved in both the TCA cycle and electron transport chain, which would be directly affected by oxygen depletion. Throughout the rest of the entire central carbon metabolism, no changes are observed.

After 10 hours of repeated oxygen oscillations, the PPP, pyruvate oxidation pathway, and TCA cycle all have several proteins in higher abundance as compared to their steady state condition. Interestingly, pyruvate oxidation and various TCA cycle reactions yield NADH, thus the increased abundance could be related to the higher NADH levels detected. The biggest effect, however, is observed in the anaerobic phase after repeated oscillations. The pyruvate oxidation, TCA cycle, and glyoxylate pathway don’t respond to the change but a sudden increase in abundance of proteins can be seen in the glycolysis and PPP. In contrast to the first anaerobic oscillation, the cells now appear to be able to quickly increase their protein content in response to anaerobic conditions. An overview of each protein’s individual profile can be found in Table S2.

These results warrant a more detailed analysis of central carbon metabolism genes. log2 fold changes of the TPM values were calculated as compared to the reference steady state (Figure 5B). Over the course of the first oscillation, no major changes occur in the transcriptome. Then, as aerobic conditions are reinstated, a small downregulation in malate synthase (YALI0_D19140g) and ribose 5-phosphate isomerase B (YALI0_F01628g) is found.

The effect of oxygen limitation remains unknown after repeated oscillations, where several genes have altered transcript levels. In glycolysis, the effect is found in the decreased expression level of two out of three hexokinases (YALI0_B22308g and YALI0_E20207g) involved in the first step of glycolysis. Interestingly, 7 proteins in glycolysis have altered abundances whereas only two genes are differentially expressed. In the PPP, ribose 5-phosphate isomerase B remains differentially downregulated which is contrasting to the 4 proteins that have an altered abundance. Meanwhile, in the TCA cycle, the gene encoding isocitrate dehydrogenase (YALI0_F04095g), involved in the conversion of iso-citrate into oxoglutarate, is upregulated. In the glyoxylate pathway conflicting behaviour is observed, as iso-citrate lyase (YALI0_C16885g) has an increased expression after exposure to repeated anaerobic oscillations while two malate synthase genes (YALI0_D19140g, YALI0_E15708g) show a downregulation.

The electron transport chain is directly linked to central carbon metabolism through its second complex, succinate dehydrogenase. Moreover, the electron transport chain is crucial as it produces NAD^+^ and ATP, and has a strict oxygen requirement. An anaerobic environment would directly affect this pathway and therefore the protein abundance in the electron transport chain was analyzed (Figure 6A). Five complexes are identified and each of them consists of a group of proteins. Similarly, to the central carbon metabolism proteins, the abundance of each protein is calculated as a log2 fold change to the steady state condition.

**Figure 6.**
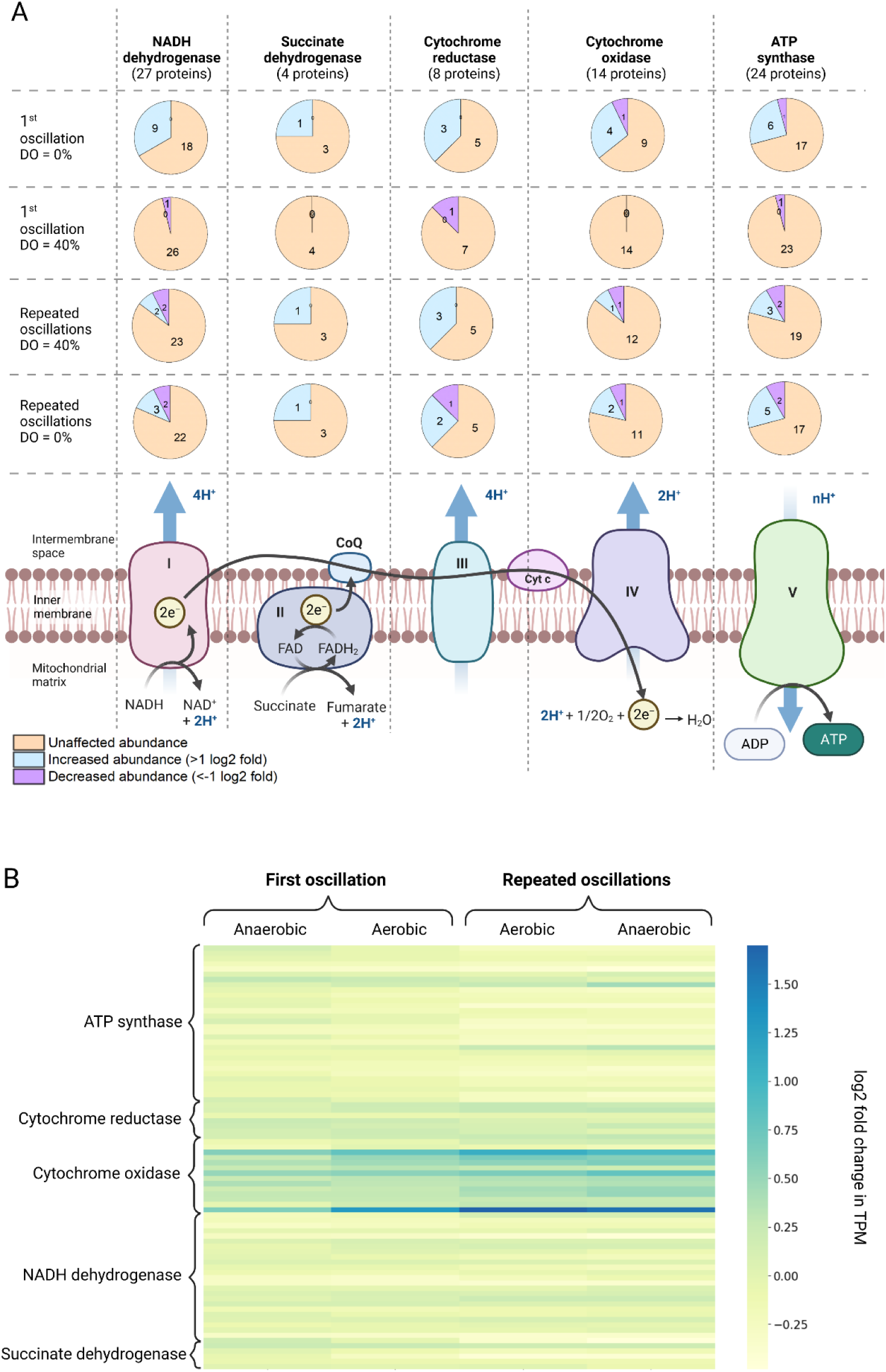
Proteomic and transcriptomic analysis of the electron transport chain. A. Changes in protein abundance in the electron transport chain over the course of a single and repeated oxygen limitations. B. Changes in TPM of genes encoding electron transport chain proteins.

The first anaerobic phase already triggers an increase in protein abundance for all electron transport chain complexes. Meanwhile, one protein in both the cytochrome oxidase and ATP synthase shows a decrease. Noteworthy is the relatively fast change in protein levels as compared to those involved in central carbon metabolism. Upon finalization of the first oscillation and a return to aerobic conditions, protein levels return to their steady state concentrations. This again indicates the relatively faster changes in the electron transport chain proteins as compared to central carbon metabolism (Table S3). Moreover, it displays an adaptability of the proteome to stimulus response experiments. Interestingly though, only two genes are found with an expression that exceeds 1 log2 fold change. The genes are both involved in cytochrome oxidase, being cytochrome C (YALI0_D09273g), and heme-a-synthase (YALI0_F24651g). Asides from that, none of the proteomic changes appear to be accompanied by differential gene expression.

After repeated oscillations, the aerobic phase shows a deviation from the values found in the steady state. At this point, there have been significant rearrangements made to the proteomic composition of the electron transport chain. When moving into the final anaerobic phase, again more proteins are affected. Interestingly, the NADH dehydrogenase complex shows a larger effect for the first oscillation as compared to the repeated one with 9 and 5 affected proteins respectively. The protein with the highest increase in abundance in cytochrome c with a log2 fold increase of 4 after repeated oxygen oscillations. Again, from a transcriptomic perspective, the only two genes with an expression that exceeds 1 log2 fold change are found in cytochrome oxidase, being cytochrome C (YALI0_D09273g), and heme-a-synthase (YALI0_F24651g).

### Anabolic shifts

Asides from the proteins involved in catabolism, it is relevant to assess the changes in anabolism. Some of the major compounds created in most microbes are composed of the building blocks that fall under the umbrella of amino acids and fatty acids. In this study, it was found that under repeated oxygen oscillations, cells increase their abundance of proteins involved with fatty acid biosynthesis (Figure 7A). More specifically, fatty acid synthase subunit alpha and enoyl-ACP reductase are both more abundant. NADPH is required for their activity, indicating an increased demand for this compound. Meanwhile, 25% of fatty acid elongation proteins show an increased abundance while fatty acid degradation pathways are affected as well. Here proteins show contradicting behavior, as there are both more and less abundant ones. The decreased abundance is found for acyl-coenzyme A oxidases and aldehyde dehydrogenase, while S-(hydroxymethyl)glutathione dehydrogenase and alcohol dehydrogenase 2 increase in abundance. Interestingly, no significant changes were found in the glycolipid pathway, which utilizes fatty acids in the production of lipids.

**Figure 7.**
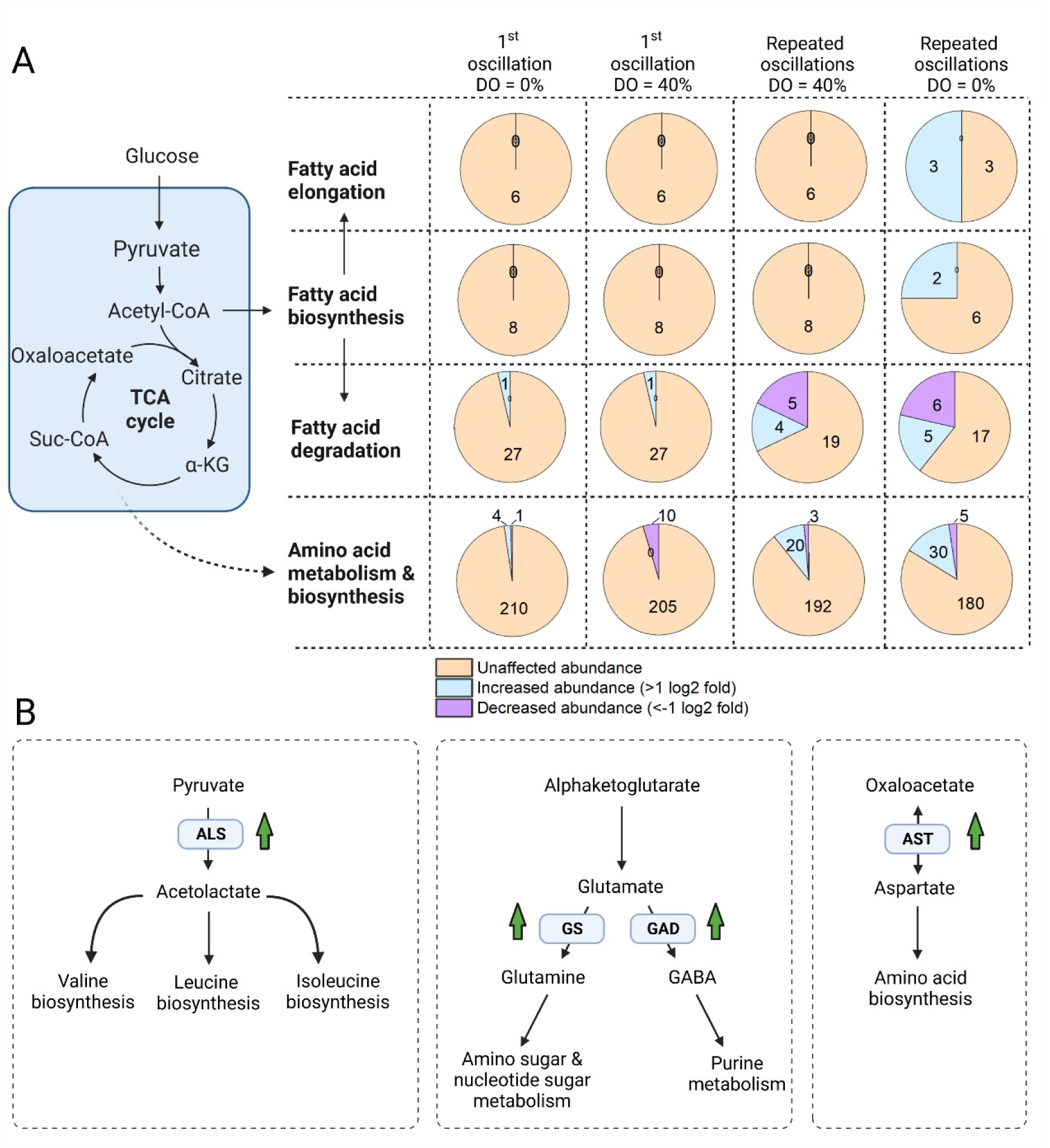
A. Proteins involved in anabolism of *Y. lipolytica*. A. Fatty acid biosynthesis directs carbon away from central carbon metabolism through Acetyl-CoA. Following this synthesis, fatty acids can be elongated or degraded. Various central carbon metabolites are also involved in the biosynthesis of amino acids. Here all proteins involved in these pathways are checked for their abundance. Samples are taken during the anaerobic phase and at the end of the first oscillation and during an aerobic and anaerobic phase 10 hours into the repeated oscillation profile. B. Increased abundance of four key proteins (ALS, GS, GAD, and AST) involved in amino acids biosynthesis after exposure to repeated oxygen oscillations. Green arrows indicated an increased abundance greater than 1 log2 fold.

On the transcriptomic side, several major changes were observed as well. One of the genes involved in fatty acid biosynthesis showed a slightly decreased expression (YALI0_E18590g), but more interestingly, a downregulation of 21 genes involved in fatty acid degradation was found (Figure S2).

Amino acid biosynthesis is a more complex process due to the wide variety of substrates and products involved. Thus, all proteins involved in general amino acid biosynthesis and metabolism were pooled together to obtain an overview of the general trends. During the first anaerobic phase, little to no changes in protein abundance are observed. After repeated oscillations on the other hand, both the aerobic and anaerobic phase show a significant increase in the abundance of proteins. Four relevant proteins that increase in abundance are acetolactate synthase (ALS), glutamate decarboxylase (GS), glutamine synthetase (GAD), and aspartate aminotransferase (AST) (Figure 7B). All four enzymes play a crucial role in amino acid biosynthesis and their increased abundance indicates an increased production of these amino acids. Moreover, all proteins are in close approximation of central carbon metabolism and thus have the potential to influence many downstream metabolite pools. Proteomic results have suggested an increased protein content after exposure to repeated anaerobic conditions. As amino acids are the key building blocks for these proteins, change in the cellular build-up would require increased amino acid production.

In line with the observations made in the proteomic analysis, the genes encoding acetolactate synthase, glutamate decarboxylase, glutamine synthetase, and aspartate aminotransferase were checked but no significant up or down-regulation was found.

## Discussion

### Steady state conditions

A chemostat was selected for this experimental setup as it allows for the determination of steady state conditions to which subsequent dynamics can be compared. The comparison with central carbon metabolite values obtained from literature shows a relatively good agreement but most notably TCA cycle intermediates citrate and succinate are much higher in this study. The metabolomics samples are taken through rapid quenching of the cells and subsequent separation of the cell pellet from the fermentation broth. The high concentration of citrate suggests that the latter step might have been executed inconsistently and considerable extracellular concentrations are measured. For the other metabolites, this effect is less obvious as these are not transported out of the cells as much. Moreover, higher standard deviations might be caused by leaking of metabolites due to sampling not happening rapidly enough, the use of suboptimal quenching solutions or inadequate temperature shocks.

Moreover, the adenylate, catabolic and anabolic redox compounds all appear to be at least twofold off from the literature values. Even though changes are expected between strains, this underlines the importance of accurate sampling. Although the obtained values remain relevant for analysis, this observation should be considered when utilizing the data for other quantitative efforts such as metabolic modelling. Meanwhile, the energy charge (Ball and Atkinson 1975; Ask et al. 2013; Suarez-Mendez et al. 2014; Czajka et al. 2020; Minden et al. 2022), catabolic (Ask et al. 2013; Minden et al. 2022) and anabolic reduction charges (Ask et al. 2013; Zhang et al. 2015; Minden et al. 2022) are all in line with values reported in literature.

For the repeated oscillating conditions, it is not likely that the cell culture grew at the designed growth rate of 0.1 h^-1^. As cells underwent anaerobic conditions, thereby decreasing ATP production and shifting catabolic and anabolic reduction charges, carbon uptake and metabolism decreased. This results in an accumulation of extracellular substrate, which is consumed once aerobic conditions are restored. Due to increased substrate levels however, cells will be able to grow at higher rates as described by Monod kinetics. As a result, the culture has a dynamic growth rate, ranging from 0 to values higher than 0.1 h^-1^, which was confirmed through compartmental modelling efforts (Kerssemakers et al. 2023). In turn, concentrations of metabolites and proteins could be affected and their comparison to the steady state value might not always be the fairest approach. An increased growth rate might for example results in an increased gene copy number, which in turn could translate to higher protein levels and altered fluxes. Moreover, an altered growth rate will result in different specific substrate uptake rates and yield factors which could potentially explain the observed differences. As it is unknown at which exact growth rate cells would be growing during the oscillations, it is difficult to correct for this influence but important to consider nonetheless.

### Cellular dynamics in response to a single oscillation

As oxygen values decrease and anaerobic conditions are established, the electron transport chain is directly affected in its fourth complex and ATP production decreases. Consequently, the glycolysis now becomes a more important contributor to the cells’ energy production. ATP is generated in the conversion of phosphoenolpyruvate to pyruvate, which could be an explanation why phosphoenolpyruvate depletes. This however is not sufficient energy to maintain a steady microbial functioning as is displayed by a rapid drop in the energy charge of the cells. As a result, glycolysis and the TCA cycle are interrupted, which affects metabolite concentrations in all pathways. Even more so, cells cannot maintain their anabolic and catabolic reduction charges, which further limits the cells’ redox potential. Interestingly, during this first anaerobic phase, the central carbon metabolism transcriptome and proteome do not show any significant changes. As shifts in the whole transcription-translation chain generally occur in a timeframe upwards of 10 minutes, which is slower than that of metabolite changes, this finding is in line with expectations (Soufi et al. 2009). In contrast, the electron transport proteome does show quick changes in response to this perturbation. Since an altered transcription was not observed, an increased post-transcriptional, translational and protein degradation regulation might be underlying this observation. It has been demonstrated that there is a significant role for these post-transcriptional regulatory events (Vogel and Marcotte 2012). This is in line with the decreasing Pearson correlation coefficient found in this work. As mRNA and protein levels do not change at the same time, and translation rates differ between proteins, a dynamic profile is to be expected (Liu et al. 2016). Moreover, it has been shown that differences in protein abundance are for only 20 to 40% caused by variable mRNA levels, stressing the impact of post-transcriptional regulation (Brockmann et al. 2007). As the Pearson correlation coefficient does not exceed 0.630, these observations support the findings of this work (Kerkhoven et al. 2017). As such it has been hypothesised that, on the short term, enzyme activity is modulated through post-translational modification and allosteric effectors (Oliveira and Sauer 2012).

Potentially relevant is the fact that mitochondria contain part of their own DNA while other parts have to be imported. This compartmentalization could have a direct effect on the speed at which cellular machinery can be adjusted. To better understand mitochondrial control, the role of AMP-activated protein kinase (AMPK) must be addressed. This molecule acts as sensor for low ATP concentrations and shows a fast response to many microbial stresses. When changes in the ATP/AMP ratio are detected, AMPK is activated which in turn phosphorylates targets to reroute metabolism towards increased catabolism and reduced anabolism. Amongst others, lipid homeostasis, glycolysis, and mitochondrial homeostasis are key targets of the AMP signaling pathway (Hedbacker and Carlson 2008; Herzig and Shaw 2017). Theoretically, it can be expected that under the selected conditions AMPK activation will occur, thereby playing a role in the observed changes.

Once aerobic conditions are restored, metabolite concentrations mostly bounce back to their original values. Also, the energy charge restores rapidly, which indicates that over a single perturbation, cells are not irreversibly damaged and are able to resume metabolic activity instantly. Moreover, the central carbon metabolism proteome remains unaffected, and the electron transport chain proteome returns to its original values. This rapid response is of great relevance as it shows a high degree of adaptability of the proteins involved in the electron transport chain. Although catabolism has shown a dynamic profile over this single oscillation, no effects were found in the anabolic parts of cellular functioning. This is an indication that no significant cellular rearrangements have occurred.

### Repeated oscillations

As the cell culture is exposed to repeated oscillations, several relevant observations were made. From both metabolomics and proteomic analysis, it appears that several metabolites, proteins, and mRNA levels differ from their steady state values after repeated oscillations. This indicates a form of adaptation or cellular rearrangement in response to constantly fluctuating conditions.

Studies on the fungi *Aspergillus fumigatus*, *Aspergillus nidulans,* and *P. pastoris* confirm these results with an increased abundance of glycolytic, TCA cycle, and PPP proteins after exposure to hypoxic conditions (Shimizu et al. 2009; Baumann et al. 2010; Vödisch et al. 2011; Barker et al. 2012). As glycolysis provides precursor molecules for cell wall biosynthesis and produces ATP, the increased protein abundance is hypothesized to have different reasons. First, as ATP generation is reduced under hypoxic conditions, upregulation of glycolysis would partially compensate for this. Secondly, hypoxic conditions were found to alter the fungal cell wall composition and thickness, thus an increased abundance of precursors could be needed (Shepardson et al. 2013; Kroll et al. 2014).

Although glycolysis, TCA cycle, and PPP showed strong dynamics with respect to the metabolome and proteome levels, the corresponding transcriptomic response was small. In glycolysis, the only adjustment was observed for two hexokinases that act as starting point of glycolysis and consequently have the potential to strongly affect all downstream activity. These findings are in line with a study on *S. cerevisiae* exposed to anaerobic conditions. Significant changes in glycolytic flues were found but these were minimally supported by transcriptional regulation, resulting in the conclusion that metabolic regulation plays a crucial role (Daran-Lapujade et al. 2007). Meanwhile, only one gene in both the TCA cycle and PPP showed an altered transcription. These results indicate that central carbon metabolism is tightly regulated at a post-transcriptional level. In line with these observations is the fast change in central carbon metabolism proteome when transitioning from the aerobic phase into the anaerobic phase. During the first perturbation, these changes were not found, indicating that over the course of repeated oscillations cells have adapted their machinery for a faster and more flexible post-transcriptional response.

The same studies also found an increase in the abundance of proteins involved in respiration and oxidative phosphorylation. Additionally, the number of mitochondria and respiratory capacity both increased under short and longer exposures to hypoxia. Interestingly not all proteins of each respiratory complex change. It is hypothesized that this could be a way to better deal with anaerobic and hypoxic conditions. It has been described that these conditions can lead to an increase in reactive oxygen species formation due to the premature transfer of electrons at complex I or III. By rearranging the complex compositions, this risk might be reduced, contributing to an increased cell health (Semenza 2007). On the other hand, ROS have also been identified to contribute to cell signalling in response to anaerobic conditions, thus controlling their levels is important for an adequate response (Guzy et al. 2007). Analysis of the transcriptome did not show any significant up- or downregulation of the genes involved in most complexes. This observation once again underlines the importance of post-transcriptional regulation, whose complexity makes it difficult to draw linear correlations between transcriptome and proteome. The only noticeable changes were found in the cytochrome oxidase complex with the largest increase in abundance of the mRNA encoding for cytochrome C. This last observation is in line with the proteomic results which showed an increased abundance of cytochrome C as well. To better understand the interactions between metabolome, proteome, and transcriptome their interactions need further studying. Relevant work has been performed in yeast where a machine learning approach is used to explore regulatory associations between metabolism and signal transduction (Gonçalves et al. 2017). Moreover, work on *S. cerevisiae* found that there is only a limited number of transcription factors (TF) that drive the change from respiratory to fermentative metabolism (Fendt et al. 2010). When exposed to different stresses the impact of post-transcriptional mechanisms was also found in yeast (Lawless et al. 2009). Moreover, post-translational modification appeared to have a strong influence on TF activity in yeast (Everett et al. 2011).

In *A. fumigatus* the proteomics analysis showed an increase in PPP proteins, which was hypothesized to be needed for an increased NAPDH demand. Indeed, several NADPH-consuming enzymes increased in abundance, among which several involved in the GABA shunt. This is in line with the increased amount of glutamate decarboxylase observed in this study. The GABA shunt bypasses two steps of the TCA cycle and is hypothesized to be involved in cell signalling in response to various stresses and acts as a possible bypass in case of TCA cycle blockage (Panagiotou et al. 2005; Cao et al. 2013). Lastly, it was also found that hypoxic conditions lead to an increased amino acid metabolism in *P. pastoris*, confirming the observation made in this work.

After repeated oscillations, central metabolite pools all decrease when transitioning from the aerobic to anaerobic phase. This is in contrast to the first stimulus-response experiment where anaerobic conditions lead to the accumulation of succinate, fumarate, and malate. However, an increased abundance of several proteins channelling carbon away from central carbon metabolism to amino acid and fatty acid production was observed as well. This potentially results in an increased flux which would lead to a depletion of central carbon metabolites. Meanwhile, this increased anabolic activity comes at the cost of an increased NADPH demand. Interestingly, the PPP metabolites and proteins both show an increased abundance after repeated oxygen oscillations. As this pathway is a primary producer of NADPH in the cells, this could be a strong indicator that flows through the PPP are increased to secure sufficient anabolic reduction power (Wasylenko et al. 2015). Transcriptional analysis of the NADPH-consuming anabolic proteins did not show any significant up- or downregulation, which underlines the complex nature of these mechanisms. Nonetheless, a large impact was seen in fatty acid metabolism through a large downregulation of a majority of genes involved. As the protome showed an increased abundance of fatty acid biosynthesis proteins, cells might respond to repeated oxygen limitation with fatty acid or lipid accumulation.

### Consequences for industrial processes with *Y. lipolytica*

Temporal anaerobic conditions are only one out of several heterogeneities that cells can experience in an industrial setting. For a strictly aerobic organism however, it is a tough one to deal with. Oxygen is a prerequisite for cellular functioning and therefore it is impossible to force the cell to maintain performance under anaerobic conditions as one could perhaps aim to do for temperature and pH fluctuations. Instead, when designing a *fermenterphile* phenotype, the main aim should be to ensure a limited burden in the loss of energy and carbon through repeated rearrangements of the cellular layout.

In this work, growth, biomass yield, and substrate uptake rate were considered as performance indicators and no specific product was considered. In an industrial setting, however, it is likely that certain production pathways are introduced or optimized. Consequently, this will alter the desired optimal cellular behaviour and changes observed in this work could be disadvantageous depending on the selected strategy. Repeated oxygen oscillations appear to increase the demand for anabolic reduction charge while anabolic reduction power struggles to keep up. Thus for most production pathways, which most likely rely on the supply of additional anabolic reduction power, this could be a problematic scenario.

This study aimed to elucidate the effects of industrially relevant oxygen oscillations on the cellular performance of *Y. lipolytica*. The anaerobic phase of a single oxygen perturbation results in dynamic metabolite pools and negatively affects the energy, catabolic and anabolic reduction charge of the cells. Metabolites either accumulate or deplete over the anaerobic phase but their concentrations recover to the original steady state values as oxygen is reintroduced. The central carbon metabolism proteome and transcriptome remain unaffected throughout the entire perturbation. In the electron transport chain, fast proteomic changes are observed but these are not accompanied by any transcriptional differences. Meanwhile, these proteomic changes revert again once aerobic conditions are reinstated. This study indicates that *Y. lipolytica* is robust and maintains a largely unaltered machinery when exposed to a single oxygen perturbation.

After 10 hours of oscillating between an industrially relevant aerobic/anaerobic profile, most metabolite concentrations differ from their steady state reference values. Whereas the energy charge and anabolic reduction charge are kept relatively constant, the catabolic reduction charge has shifted significantly from its steady state value. This indicates a shift in redox homeostasis and a loss in catabolic reduction power. Moreover, proteomic results indicate an increased catabolic activity related to amino acid and fatty acid biosynthesis. This would come at the cost of additional NADPH demand which is likely to be provided by increased PPP activity. When shifting from aerobic to anaerobic conditions, the proteome now quickly adjusts itself as compared to the initial perturbation. The relatively fast changes in the proteome when transitioning from aerobic to anaerobic conditions, alongside the limited changes to the associated transcriptome, lead to the conclusion that post-transcriptional regulation plays a major role in *Y. lipolytica* metabolism and adaptation to oxygen oscillations. Moreover, these findings showcase that with this altered transcriptomic, proteomic and metabolomics layout, *Y. lipolytica* can maintain almost comparable growth performance under repeated anaerobic conditions.

### Data availability

All RNA-Seq. data has been uploaded to the Sequence Read Archive (SRA) and is further described under BioProject number PRJNA947955. Proteomics and metabolomics data is available in the supplementary material of this manuscript.

## Acknowledgements

We thank the Yeast Metabolic Engineering group at the Novo Nordisk Foundation Center for Biosustainability for providing the W29 strain used in this research. Some figure in this work were (partially) created through Biorender.

## Author contributions

Conceptualization, S.S. and A.A.J.K.; methodology, S.S., M.N.F, S.Ø. and A.A.J.K.; software, S.Ø., M.N.F and A.A.J.K.; validation, M.N.F., S.Ø. and A.A.J.K.; formal analysis, S.Ø., M.N.F. and A.A.J.K.; investigation, S.S., M.N.F., S.Ø. and A.A.J.K.; resources, S.S.; data curation, A.A.J.K.; writing—original draft preparation, A.A.J.K.; writing—review and editing, S.S., M.N.K., S.Ø. and A.A.J.K.; visualization, A.A.J.K.; supervision, S.S.; project administration, S.S.; funding acquisition, S.S. and A.A.J.K. All authors have read and agreed to the published version of the manuscript.

## Compliance with Ethical Standards

• All authors declare that he/she has no conflict of interest.

• This article does not contain any studies with human participants or animals performed by any of the authors.

