## Supplementary material for "The dominant role of post-transcriptional regulation in *Yarrowia lipolytica* to repeated O_2_ limitations"

### Supplementary Table 1 – Steady state metabolite concentrations

Table 1. Steady state metabolite concentrations across varying yeast species. The literature values on *S. cerevisae* and *Y. lipolytica* are estimated based on plots from their respective publications.

| **Metabolite** | **Steady state value (µmol/gCDW)** | ***S. cerevisiae*** (Minden et al. 2022) **(µmol/gCDW)** | ***Y. lipolytica*** (Christen and Sauer) **(µmol/gCDW)** | ***P. pastoris*** (Carnicer et al. 2012) **(µmol/gCDW)** |
|---|---|---|---|---|
| G6P | *6.90 ± 5.48* | *8* | *3* | *6.4* |
| F6P | *5.31 ± 3.45* | | *0.5* | *1.4* |
| G3P | *1.48 ± 0.49* | | | *2.1* |
| PEP | *0.66 ± 0.16* | *2* | *0.05* | *0.48* |
| AcCoA | *3.86 ± 0.25* | | | |
| CIT | *8715.17 ± 4900* | | | *10.7* |
| AKG | *0.87 ± 0.11* | *1,8* | *3.5* | *2.9* |
| SUCC | *18.47 ± 2.77* | *0.8* | *0.5* | *3.3* |
| FUM | *0.49 ± 0.05* | *0.5* | | *1.3* |
| MAL | *5.69 ± 0.80* | *9* | | *7.2* |
| RU5P | *0.14* | | | |
| R5P | *0.35 ± 0.09* | | *0.4* | |
| AMP | *0.83 ± 0.37* | *0.25* | *0.2* | |
| ADP | *2.80 ± 0.23* | *2* | *0.8* | |
| ATP | *19.89 ± 3.57* | *9* | *5* | |
| NAD+ | *8.36 ± 3.45* | *4* | | |
| NADH | *0.88 ± 0.44* | *0.17* | | |
| NADP+ | *1.57 ± 0.25* | *0.15* | | |
| NADPH | *1.36 ± 0.53* | *0.2* | | |

Supplementary Figure 1 - Pearson correlation coefficients

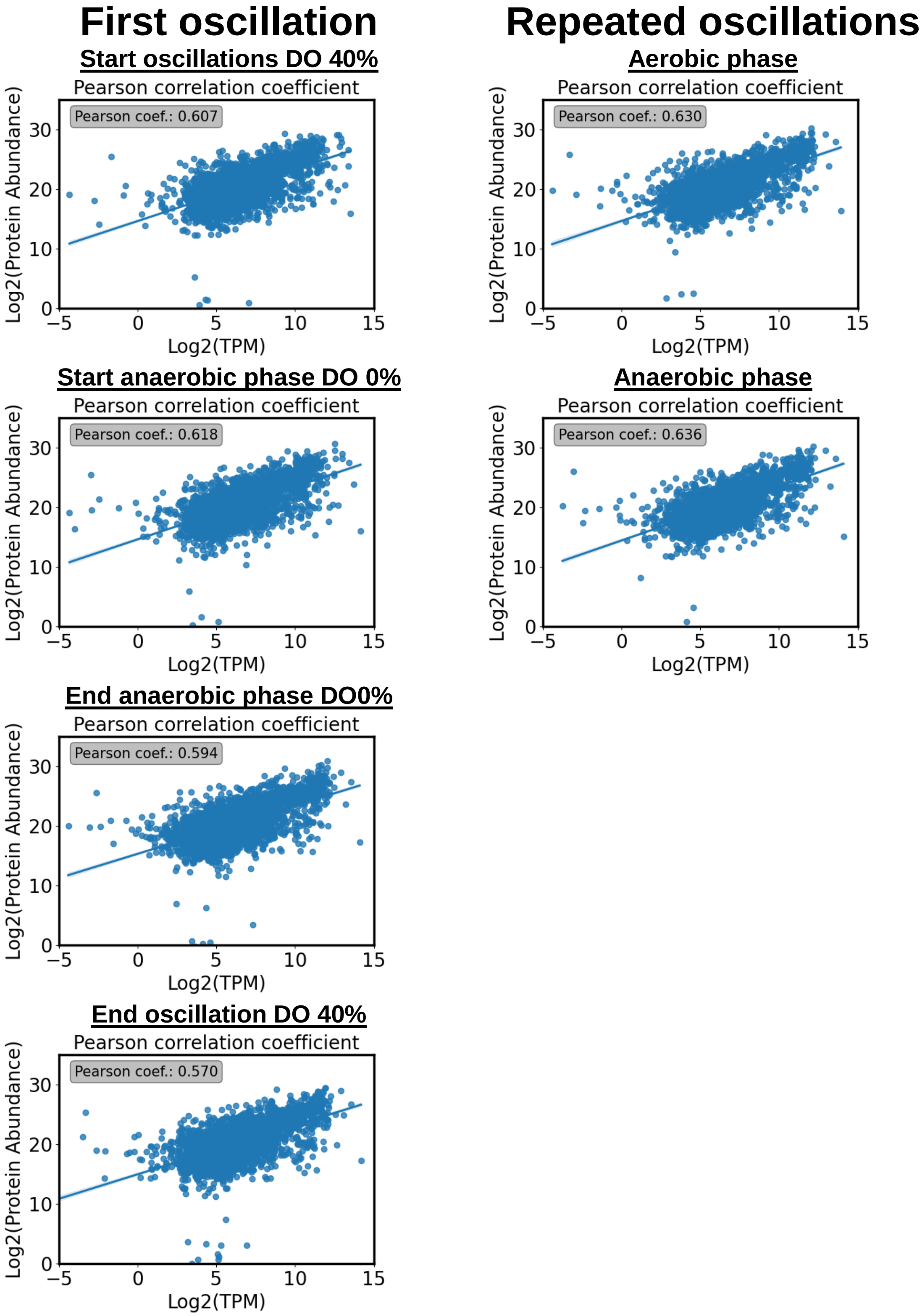

### Supplementary Table 2- Central carbon metabolism proteins

| UniProt ID | Enzyme | Pathway | Anaerobic | Aerobic | Aerobic | Anaerobic |
|---|---|---|---|---|---|---|
| F2Z672 | Phosphotransferase | Glycolysis | 0.404455 | 0.519082 | 0.651939 | 0.673383 |
| P29407 | Phosphoglycerate kinase | Glycolysis | 0.499032 | 0.353644 | 0.927253 | 1.208305 |
| P30614 | Pyruvate kinase | Glycolysis | 0.35319 | -0.1282 | 0.961942 | 1.371733 |
| P59680 | ATP-dependent 6-phosphofructokinase | Glycolysis | 0.229148 | 0.354356 | 0.725051 | 1.065868 |
| Q6C1F3 | Enolase | Glycolysis | 0.352208 | 0.098214 | 0.650759 | 1.019265 |
| Q6C2I1 | Glucose-6-phosphate isomerase | Glycolysis | 0.108843 | 0.274427 | 0.277465 | 0.430753 |
| Q6C4K5 | Fructose-bisphosphate aldolase | Glycolysis | 0.399468 | 0.20104 | 0.680835 | 1.016403 |
| Q6C5S9 | Phosphotransferase | Glycolysis | -0.18547 | 0.108895 | -0.04744 | -0.01249 |
| Q6CCU7 | Glyceraldehyde-3-phosphate dehydrogenase | Glycolysis | 0.374408 | 0.35424 | 0.663077 | 1.049814 |
| Q6CFX7 | Phosphoglycerate mutase | Glycolysis | 0.781562 | -0.1526 | 0.790763 | 1.099435 |
| Q6CGV1 | fructose-bisphosphatase | Glycolysis | 0.113437 | 0.213719 | 0.163711 | 0.220386 |
| P41555 | Isocitrate lyase | Glyoxylate pathway | 0.092262 | 0.141031 | 0.624906 | 0.814697 |
| Q6BZP5 | 2-methylisocitrate lyase, mitochondrial | Glyoxylate pathway | -0.28976 | 0.337228 | -0.046 | 0.019344 |
| Q6C5R9 | Malate synthase | Glyoxylate pathway | 0.338342 | 0.273836 | 0.125379 | 0.367999 |
| Q6C8J4 | Malate synthase | Glyoxylate pathway | -1.54768 | -1.27286 | -0.89442 | 0.082878 |
| Q6C5F0 | YALI0E18634p | Other | 0.242627 | 0.595748 | 1.777871 | 1.970308 |
| Q6CAV2 | Pyruvate carboxylase | Other | 0.072972 | 0.340306 | 0.527752 | 0.624309 |
| Q6C1K2 | Transaldolase | Pentose phosphate pathway (non-ox) | -0.04537 | 0.244636 | 0.35526 | 0.415579 |
| Q6C2T9 | Triosephosphate isomerase | Pentose phosphate pathway (non-ox) | 0.48606 | 0.19321 | 0.491187 | 1.132518 |
| Q6C393 | YALI0F01628p | Pentose phosphate pathway (non-ox) | 0.18478 | 0.084813 | 0.727003 | 0.847727 |
| Q6C4K5 | Fructose-bisphosphate aldolase | Pentose phosphate pathway (non-ox) | 0.399468 | 0.20104 | 0.680835 | 1.016403 |
| Q6C6T4 | Transketolase | Pentose phosphate pathway (non-ox) | 0.456658 | 0.576127 | 0.918088 | 1.250772 |
| Q6CAJ3 | YALI0D02277p | Pentose phosphate pathway (non-ox) | 0.176216 | 0.556945 | 0.687443 | 0.750681 |
| Q6CC71 | Ribulose-phosphate 3-epimerase | Pentose phosphate pathway (non-ox) | -0.56439 | -0.14906 | -0.18522 | -0.0337 |
| Q6CFH4 | Ribose-5-phosphate isomerase | Pentose phosphate pathway (non-ox) | -0.03664 | 0.107025 | -0.26639 | -0.3376 |
| Q6C4Y7 | Glucose-6-phosphate 1-dehydrogenase | Pentose phosphate pathway (ox) | -0.05754 | 0.099312 | 0.168607 | 0.235838 |
| Q6C682 | 6-phosphogluconolactonase-like protein | Pentose phosphate pathway (ox) | -0.06674 | -0.00899 | -0.38046 | -0.32134 |
| Q6CBG4 | 6-phosphogluconolactonase-like protein | Pentose phosphate pathway (ox) | 0.875176 | 0.340596 | 1.349027 | 1.455539 |
| Q6CEH4 | 6-phosphogluconate dehydrogenase, decarboxylating | Pentose phosphate pathway (ox) | 0.08567 | 0.190676 | 0.487701 | 0.761966 |
| Q6C0Y7 | Pyruvate dehydrogenase E1 component subunit alpha | Pyruvate oxidation | 0.178781 | 0.11498 | 0.340583 | 0.581078 |
| Q6C4G4 | Pyruvate dehydrogenase E1 component subunit beta | Pyruvate oxidation | 0.717498 | -0.2658 | 1.371849 | 1.774895 |

| | | | | | | |
|---|---|---|---|---|---|---|
| Q6C812 | Acetyltransferase component of pyruvate dehydrogenase complex | Pyruvate oxidation | 0.393643 | 0.313289 | 0.625514 | 0.696266 |
| Q6C8C6 | Dihydrolipoyl dehydrogenase | Pyruvate oxidation | 0.695763 | 0.188462 | 1.0761 | 1.397936 |
| Q6C2Y4 | Isocitrate dehydrogenase [NADP] | TCA cycle | 0.059164 | 0.129388 | 0.254278 | 0.322024 |
| Q6C3M8 | oxoglutarate dehydrogenase (succinyl-transferring) | TCA cycle | 0.214858 | 0.134354 | 0.492205 | 0.83489 |
| Q6C450 | YALI0E29667p | TCA cycle | #VALUE! | #VALUE! | #VALUE! | #VALUE! |
| Q6C4S9 | Succinate--CoA ligase [ADP-forming] subunit alpha, mitochondrial | TCA cycle | 0.02853 | 0.262784 | 0.283708 | 0.578596 |
| Q6C5L8 | dihydrolipoyllysine-residue succinyltransferase | TCA cycle | 0.0777 | 0.159896 | 0.19223 | 0.209923 |
| Q6C5X9 | malate dehydrogenase | TCA cycle | -0.01078 | 0.187885 | 0.566774 | 0.587793 |
| Q6C6Z1 | isocitrate dehydrogenase (NAD(+)) | TCA cycle | 0.174986 | 0.28256 | 0.474111 | 0.552342 |
| Q6C793 | 2-methylcitrate synthase, mitochondrial | TCA cycle | 0.432643 | 0.310659 | 0.666 | 0.939494 |
| Q6C7I2 | Citrate synthase | TCA cycle | -0.15285 | 0.212972 | 0.376505 | 0.516002 |
| Q6C823 | Succinate dehydrogenase [ubiquinone] iron-sulfur subunit, mitochondrial | TCA cycle | -0.05355 | -0.09527 | 0.015614 | 0.12273 |
| Q6C8C6 | Dihydrolipoyl dehydrogenase | TCA cycle | 0.695763 | 0.188462 | 1.0761 | 1.397936 |
| Q6C8V3 | Malate dehydrogenase | TCA cycle | -0.19875 | -0.23645 | 0.062798 | 0.207253 |
| Q6C9G6 | Succinate dehydrogenase [ubiquinone] flavoprotein subunit, mitochondrial | TCA cycle | 0.624232 | 0.476714 | 0.757201 | 0.915397 |
| Q6C9P6 | Aconitate hydratase, mitochondrial | TCA cycle | 0.118128 | 0.188268 | 0.329562 | 0.499427 |
| Q6CA33 | isocitrate dehydrogenase (NAD(+)) | TCA cycle | -0.23941 | -0.03303 | 0.010385 | 0.109004 |
| Q6CA97 | Succinate--CoA ligase [ADP-forming] subunit beta, mitochondrial | TCA cycle | -0.1455 | -0.01505 | 0.25076 | 0.268417 |
| Q6CCT2 | fumarate hydratase | TCA cycle | 0.147506 | 0.38235 | 0.531978 | 0.835439 |
| Q6CGZ0 | Succinate dehydrogenase [ubiquinone] cytochrome b small subunit | TCA cycle | 2.077497 | 0.611376 | 1.360958 | 1.920094 |

### Supplementary Table 3 - Electron transport chain proteins

| UniProt ID | Enzyme | Module | Anaerobic | Aerobic | Aerobic | Anaerobic |
|---|---|---|---|---|---|---|
| B5FVB3 | ATP synthase subunit K, mitochondrial | ATP port | 0.957861 | -0.15505 | 1.001132 | 1.565865 |
| B5FVB8 | ATP synthase subunit g, mitochondrial | ATP port | -0.55386 | -0.96987 | -1.02721 | -1.23751 |
| B5FVG3 | ATP synthase subunit e, mitochondrial | ATP port | 1.164021 | 0.135968 | 1.225958 | 1.629483 |
| Q6BZT1 | V-type proton ATPase subunit a | ATP port | 0.07802 | 0.213065 | 0.065698 | -0.04665 |
| Q6C105 | ATP synthase subunit 4, mitochondrial | ATP port | 0.409471 | 0.684094 | 0.770259 | 0.87398 |
| Q6C1H5 | V-type proton ATPase subunit | ATP port | 0.05772 | 0.058978 | -0.1381 | -0.112 |
| Q6C1K0 | YALI0F15631p | ATP port | -0.0971 | -0.23996 | -1.15394 | -1.23233 |
| Q6C1T4 | Inorganic pyrophosphatase | ATP port | -0.08306 | 0.149457 | -0.02272 | 0.103691 |
| Q6C2V6 | ATP synthase subunit H, mitochondrial | ATP port | -0.37414 | -0.73933 | -0.65967 | -0.8241 |
| Q6C326 | ATP synthase subunit alpha, mitochondrial | ATP port | 0.152754 | -0.10073 | 0.522552 | 0.695312 |
| Q6C338 | ATP synthase subunit gamma, mitochondrial | ATP port | -0.14679 | 0.030253 | -0.12916 | 0.031872 |
| Q6C4E9 | Vacuolar proton pump subunit B | ATP port | 0.020449 | 0.281247 | 0.064351 | 0.101829 |
| Q6C5Q2 | V-type proton ATPase subunit F | ATP port | 0.184639 | -0.1702 | 0.312961 | 1.160545 |
| Q6C6F6 | V-type proton ATPase subunit a | ATP port | -6.14547 | #VALUE! | -0.84566 | -0.44597 |
| Q6C6K9 | V-type proton ATPase subunit H | ATP port | -0.35378 | -0.0924 | -0.42035 | -0.18725 |
| Q6C877 | ATP synthase subunit delta, mitochondrial | ATP port | 0.763444 | -1.30033 | 0.931253 | 1.499308 |
| Q6C8S0 | ATP synthase subunit J, mitochondrial | ATP port | 1.476749 | -0.29509 | 1.353739 | 1.565694 |
| Q6C9B1 | ATP synthase subunit 5, mitochondrial | ATP port | 0.220545 | -0.22195 | -0.50987 | -0.57571 |
| Q6C9E6 | ATP synthase subunit f, mitochondrial | ATP port | 0.083263 | -0.16752 | 0.557574 | 0.387476 |
| Q6CAR5 | YALI0D00583p | ATP port | 0.535378 | 0.62026 | -0.17927 | 0.175555 |
| Q6CC75 | inorganic diphosphatase | ATP port | -0.1381 | -0.71977 | 0.45002 | 0.759438 |
| Q6CDQ7 | Plasma membrane ATPase | ATP port | 0.472236 | 0.171754 | 0.683962 | 0.487697 |
| Q6CFH9 | ATP synthase subunit d, mitochondrial | ATP port | 1.238654 | -0.47147 | 0.548478 | 0.702144 |
| Q6CFT7 | ATP synthase subunit beta, mitochondrial | ATP port | 0.28421 | 0.067131 | 0.657273 | 0.814607 |
| Q6CH91 | V-type proton ATPase subunit C | ATP port | 0.121405 | 0.173539 | 0.18225 | 0.270682 |
| Q6CHD8 | H(+)-transporting two-sector ATPase | ATP port | 0.134848 | 0.300764 | 0.269025 | 0.363873 |
| W0TYM5 | V-type proton ATPase subunit G | ATP port | 0.533136 | -0.62781 | 0.329903 | 0.625546 |
| Q6C0H4 | YALI0F24673p | Cytochrome bc1 complex | 1.361443 | #VALUE! | 1.629717 | 3.323401 |
| Q6C2E3 | Cytochrome b-c1 complex subunit 2, mitochondrial | Cytochrome bc1 complex | 0.005768 | 0.111649 | 0.438254 | 0.576237 |
| Q6C387 | Cytochrome b-c1 complex subunit 8 | Cytochrome bc1 complex | 1.408892 | -0.21892 | 1.290514 | 0.969346 |

| | | | | | | |
|---|---|---|---|---|---|---|
| Q6C3K7 | Cytochrome b-c1 complex subunit 7 | Cytochrome bc1 complex | 0.517884 | 0.151596 | 0.209929 | 0.410807 |
| Q6CC60 | YALI0C12210p | Cytochrome bc1 complex | 1.769544 | -1.34799 | 1.446077 | 1.561891 |
| Q6CG23 | Complex III subunit 9 | Cytochrome bc1 complex | -0.15699 | -0.95213 | -0.94495 | -1.05826 |
| Q6CGP7 | YALI0A17468p | Cytochrome bc1 complex | 0.239569 | 0.257065 | 0.738833 | 0.948466 |
| Q6CI02 | Cytochrome b-c1 complex subunit Rieske, mitochondrial | Cytochrome bc1 complex | -0.03375 | -0.06684 | -0.4415 | -0.30891 |
| B5FVH7 | YALI0E12628p | Cytochrome c oxidase | -0.00974 | -0.29199 | -0.63091 | -0.79602 |
| B5RSK7 | YALI0F04114p | Cytochrome c oxidase | 0.265562 | -0.82081 | 0.477846 | 0.089156 |
| Q6C094 | YALI0F26675p | Cytochrome c oxidase | -0.07447 | -0.19472 | -1.14009 | -1.17737 |
| Q6C0H5 | YALI0F24651p | Cytochrome c oxidase | 0.553995 | 0.315786 | 0.216232 | 0.153064 |
| Q6C309 | YALI0F03567p | Cytochrome c oxidase | 0.70439 | 0.221264 | 0.573848 | 0.809255 |
| Q6C325 | Cytochrome c oxidase subunit | Cytochrome c oxidase | 0.472486 | -0.0663 | 0.253767 | 0.215755 |
| Q6C5A3 | YALI0E19723p | Cytochrome c oxidase | 0.648704 | 0.639417 | 0.654726 | 0.943027 |
| Q6C5M8 | Cytochrome c oxidase subunit | Cytochrome c oxidase | 0.201799 | 0.296288 | -0.39478 | -0.30759 |
| Q6C6E6 | Cytochrome c oxidase subunit 6, mitochondrial | Cytochrome c oxidase | 0.418456 | 0.14047 | 0.768116 | 0.910767 |
| Q6C6F0 | YALI0E10065p | Cytochrome c oxidase | #VALUE! | 0.137616 | #VALUE! | 4.94516 |
| Q6C9Q0 | Cytochrome c | Cytochrome c oxidase | 2.111585 | -0.35187 | 2.73282 | 4.050314 |
| Q6CBM0 | YALI0C17391p | Cytochrome c oxidase | 0.588392 | 0.627495 | -0.7417 | -2.55812 |
| Q9B6D5 | Cytochrome c oxidase subunit 2 | Cytochrome c oxidase | 0.586451 | 0.45982 | 0.614312 | 0.581439 |
| Q9B6E4 | COX1-i3 protein, alternatively spliced | Cytochrome c oxidase | -0.10328 | #VALUE! | 0.376242 | 0.770211 |
| B5FVE5 | YALI0D10274p | NADH dehydrogenase | 0.618631 | 0.211677 | 0.678593 | 0.915812 |
| B5FVF8 | NADH dehydrogenase [ubiquinone] 1 alpha subcomplex subunit 13 | NADH dehydrogenase | 0.577883 | 0.42147 | -0.20483 | 0.148414 |
| B5FVG1 | NADH dehydrogenase [ubiquinone] 1 beta subcomplex subunit 7 | NADH dehydrogenase | 0.265448 | -0.13852 | -0.09499 | -0.02214 |
| Q6C4W9 | YALI0E23089p | NADH dehydrogenase | 0.69796 | -0.22893 | 0.628666 | 0.736468 |
| Q6C7X2 | Acyl carrier protein | NADH dehydrogenase | 0.082653 | -0.27916 | -0.44661 | -0.49594 |
| Q6C7X4 | YALI0D24585p | NADH dehydrogenase | 0.052637 | 0.424003 | 0.4003 | 0.686411 |
| Q6C926 | Acyl carrier protein | NADH dehydrogenase | 0.207738 | 0.009163 | -0.44822 | -0.48113 |
| Q6C9Z1 | NADH dehydrogenase [ubiquinone] 1 beta subcomplex subunit 9 | NADH dehydrogenase | 0.439287 | 0.075686 | -0.17497 | -0.17587 |
| Q6CA88 | YALI0D04939p | NADH dehydrogenase | 0.607276 | -1.2998 | 0.789942 | 1.402333 |
| Q6CD73 | YALI0C03201p | NADH dehydrogenase | 0.516276 | 0.149291 | 0.35216 | 0.351714 |
| Q6CG53 | NADH dehydrogenase [ubiquinone] 1 alpha subcomplex subunit N7BM | NADH dehydrogenase | -0.36766 | -0.47621 | -1.40344 | -1.38333 |
| Q6CGB4 | NADH-ubiquinone oxidoreductase | NADH dehydrogenase | 0.867602 | -0.43264 | -0.00138 | 0.446863 |
| Q6CI60 | YALI0A01419p | NADH dehydrogenase | 0.977337 | -0.18329 | 0.521014 | 0.967308 |
| B5RSL7 | YALI0F18359p | NADH dehydrogenase (ubiquinone) Fe-S | 0.888375 | 0.704837 | 0.316368 | 0.407011 |

| | | | | | | |
|---|---|---|---|---|---|---|
| | | protein/flavoprotein complex_ mitochondria | | | | |
| F2Z619 | YALI0F00924p | NADH dehydrogenase (ubiquinone) Fe-S protein/flavoprotein complex_ mitochondria | -0.4198 | -0.47452 | -1.00753 | -1.40729 |
| F2Z626 | YALI0F17248p | NADH dehydrogenase (ubiquinone) Fe-S protein/flavoprotein complex_ mitochondria | -0.20473 | 0.181739 | 0.417895 | 0.442729 |
| F2Z660 | NADH dehydrogenase [ubiquinone] flavoprotein 1, mitochondrial | NADH dehydrogenase (ubiquinone) Fe-S protein/flavoprotein complex_ mitochondria | 0.272161 | 0.193881 | 0.631656 | 0.78842 |
| F2Z6C0 | YALI0D00737p | NADH dehydrogenase (ubiquinone) Fe-S protein/flavoprotein complex_ mitochondria | -0.44767 | -0.45112 | -0.66364 | -0.50628 |
| F2Z6D7 | YALI0F02123p | NADH dehydrogenase (ubiquinone) Fe-S protein/flavoprotein complex_ mitochondria | -0.17769 | 0.176402 | 0.179396 | 0.669382 |
| F2Z6F1 | YALI0D05467p | NADH dehydrogenase (ubiquinone) Fe-S protein/flavoprotein complex_ mitochondria | 0.060904 | 0.28903 | 0.37146 | 0.489691 |
| Q6C2Q1 | YALI0F06050p | NADH dehydrogenase (ubiquinone) Fe-S protein/flavoprotein complex_ mitochondria | 0.250799 | -0.21202 | -0.31728 | -0.18677 |
| Q6C8J9 | YALI0D19030p | NADH dehydrogenase (ubiquinone) Fe-S protein/flavoprotein complex_ mitochondria | 1.405165 | -0.7204 | 1.305054 | 1.869849 |
| Q6CEK9 | NADH dehydrogenase [ubiquinone] iron-sulfur protein 4, mitochondrial | NADH dehydrogenase (ubiquinone) Fe-S protein/flavoprotein complex_ mitochondria | 0.441014 | 0.23264 | -0.25127 | -0.21281 |
| F2Z699 | External alternative NADH-ubiquinone oxidoreductase, mitochondrial | NADH:quinone reductase (non-electrogenic) | 0.169015 | 0.20413 | 0.786257 | 0.97176 |
| Q6C6X0 | YALI0E05599p | NADH:quinone reductase (non-electrogenic) | 0.296276 | 0.002188 | 0.312691 | 0.411155 |
| Q9B6D3 | NADH-ubiquinone oxidoreductase chain 5 | NADH:ubiquinone oxidoreductase_ mitochondria | -0.39751 | 0.073647 | 0.178576 | -0.07933 |
| Q9B6E8 | NADH-ubiquinone oxidoreductase chain 1 | NADH:ubiquinone oxidoreductase_ mitochondria | 0.103038 | 0.493851 | 1.748502 | 1.78812 |
| Q6C450 | YALI0E29667p | Succinate dehydrogenase (ubiquinone | #VALUE! | #VALUE! | #VALUE! | #VALUE! |
| Q6CGZ0 | Succinate dehydrogenase [ubiquinone] cytochrome b small subunit | Succinate dehydrogenase (ubiquinone | 2.077497 | 0.611376 | 1.360958 | 1.920094 |

| | | | | | | |
|---|---|---|---|---|---|---|
| Q6C823 | Succinate dehydrogenase [ubiquinone] iron-sulfur subunit, mitochondrial | Succinate dehydrogenase (ubiquinone) flavoprotein subunit | -0.05355 | -0.09527 | 0.015614 | 0.12273 |
| Q6C9G6 | Succinate dehydrogenase [ubiquinone] flavoprotein subunit, mitochondrial | Succinate dehydrogenase (ubiquinone) flavoprotein subunit | 0.624232 | 0.476714 | 0.757201 | 0.915397 |

### Supplementary Figure 2- Transcriptome analysis of fatty acid metabolism

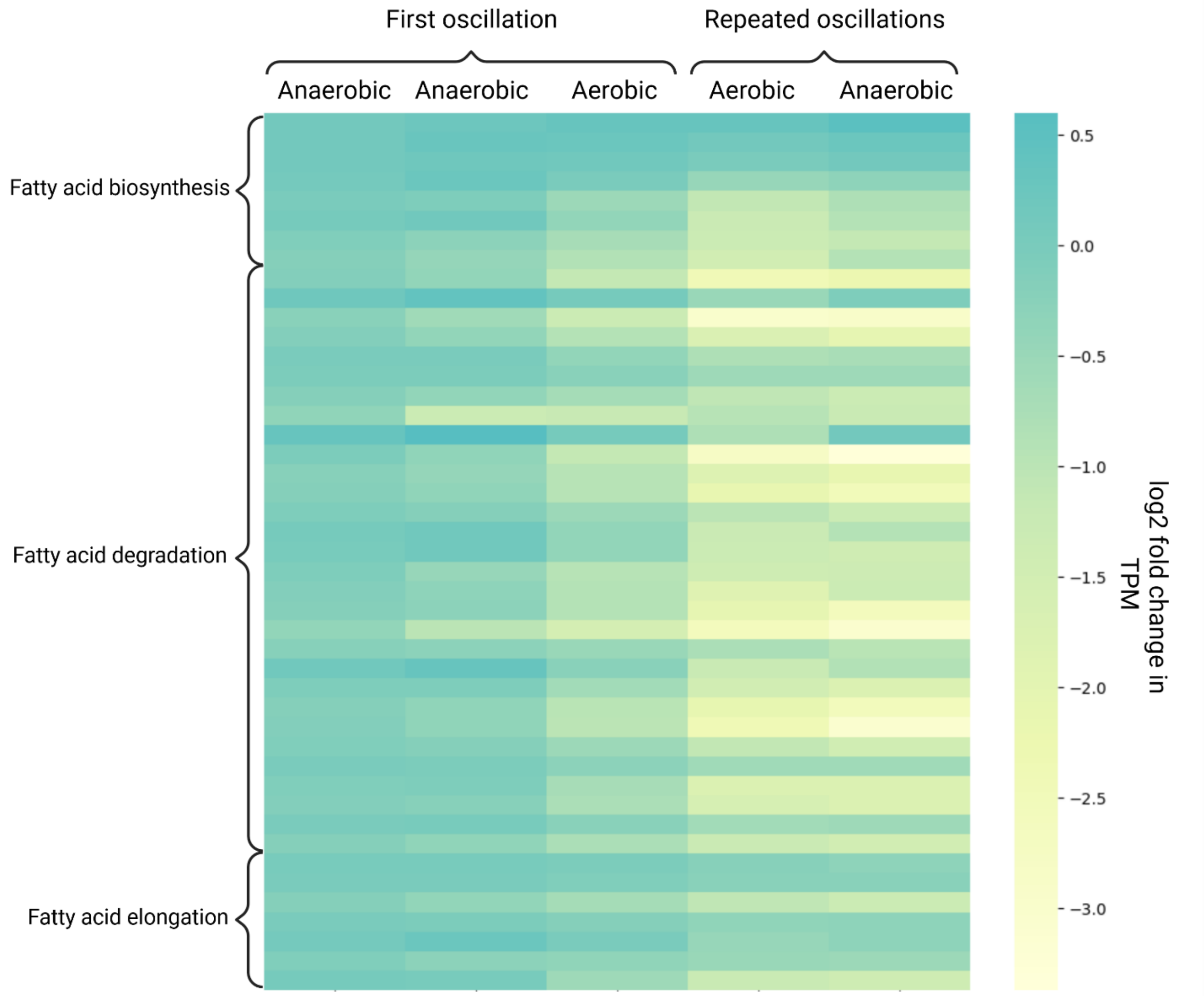
